# *Enterococcus faecium* MBBL3 Exhibits Promising Probiotic Potential and Antimicrobial Efficacy Against Bovine Mastitis-Associated *Escherichia coli and Klebsiella pneumoniae*

**DOI:** 10.1101/2025.01.07.631658

**Authors:** Naim Siddique, Md. Morshedur Rahman, Soharth Hasnat, ANM Aminoor Rahman, Anup Kumar Talukder, Md. Robiul Karim, Ziban Chandra Das, Tofazzal Islam, M. Nazmul Hoque

## Abstract

*Enterococcus faecium*, a promising probiotic, combats pathogens, supports gut health, strengthens immunity, and provides a natural approach to address the escalating global challenge of antimicrobial resistance. This study aimed to investigate the genome of *E. faecium* MBBL3, isolated from healthy cow milk, to assess its probiotic potential and antimicrobial activity against pathogens causing bovine mastitis. The strain was analyzed through whole genome sequencing, along with *in-vitro* and *in-silico* assessments were conducted to determine its antimicrobial efficacy against bovine mastitis pathogens, *Klebsiella pneumoniae* MBBL2 (*Kp* MBBL2) and *Escherichia coli* MBBL4 (*Ec* MBBL4). The genome assembly and functional annotations uncovered many important probiotic traits in MBBL3, where genome comparison revealed its high genetic similarity with other *Enterococcus* strains. MBBL3 demonstrated the ability to ferment a wide range of carbohydrates and possessed 76 carbohydrate-active enzyme-related genes, including five key CAZy families namely GH73, GH18, CBM50, CE4, and AA10. It also possessed importance genes for bile salt and acid tolerance, stress resistance, and surface adhesion. Additionally, MBBL3 contained metabolite regions involved in the biosynthesis of antimicrobial compounds such as 2,4-DAPG, aborycin, enterocin NKR-5-3B, and sodorifen, and bacteriocin gene clusters for *sactipeptides*, *Enterolysin_A*, and *UviB*. Safety assessments indicated low pathogenic potential, while *in-vitro* assays demonstrated antibiotic susceptibility and suppressed the growth of *Kp* MBBL2 and *Ec* MBBL4, respectively. Its bacteriocin compound Enterolysin_A exhibited strong molecular interactions with virulence proteins of these mastitis pathogens. Therefore, the promising probiotic potential and antimicrobial efficacy of *E. faecium* MBBL3, especially against mastitis pathogens combined with its safety, position it as a valuable candidate for therapeutic applications.

**Key points:**

- *E. faecium* MBBL3 genosme showed high similarity with other species of this genera.
- Genetic makeup of MBBL3 revealed its ability to survival and adaptation in different niches including host’s gut.
- *In-vitro* and *in-silico* study results, along with several genes linked to antimicrobials demonstrated its ability to combat against mastitis pathogens.

## Introduction

Probiotics are known as the living microorganisms which provide beneficial impact on the host health (Chugh and Kamal-Eldin 2020). In recent years, probiotics have emerged as a focal point of scientific research due to their profound impact on host health. Among the diverse range of probiotic bacteria, *Enterococci* have garnered significant scientific interest for their potential health benefits, including antimicrobial activity, gut microbiota modulation, immune enhancement, bioactive compound production, promoting gastrointestinal and overall health (Ben Braïek and Smaoui 2019; Qiu et al. 2022). *E. faecium* is a major species of the genus *Enterococcus*, that can thrive in various niches due to their adaptive features to survive in a variety of environments enabling them to colonize in diverse hosts and niches (Ness et al. 2014). They are naturally found in fermented food items as well as gastrointestinal tract of human and other animals (Hanchi et al. 2018). This microorganism was found to play a vital role in the naturally fermented foods, enhancing their texture, flavors and aroma by producing various aromatic compounds, enzymes and different metabolites (Žugić Petrović et al. 2020). Additionally, *E. faecium* exhibits wide range of metabolic capabilities, including carbohydrate and amino acid fermentation, and synthesizes essential vitamins which enhance its acceptance in the food industry (Aswal et al. 2023). It has also been reported that *E. faecium* play beneficial role in modulation of gut microbiota, inhibition of pathogen adhesion, while increasing the gastrointestinal barrier function, inducing anti-inflammatory cytokine production and suppressing pro-inflammatory cytokines, as well as exhibiting the cholesterol-lowering activity (Holzapfel et al. 2018; Zhang et al. 2017). *E. faecium* is well-recognized for producing natural antimicrobials such as antimicrobial peptides (AMPs) and bacteriocins being a potential probiotic bacterium (Tedim et al. 2024; Zommiti et al. 2018). The beneficial effects of *E. faecium* on immune modulation and its capacity to inhibit pathogenic bacterial growth have been supported by both *in-vitro* and *in-vivo* studies (Lodemann et al. 2015). The impact of its AMPs and bacteriocins on the colonization of pathogenic bacteria, including *Escherichia coli* and *Salmonella enteritidis*, has been extensively documented (Bhardwaj et al. 2010; Palkovicsné Pézsa et al. 2022). Additionally, probiotic strains of *E. faecium* have been shown to promote the growth of beneficial bacteria, such as *Lactobacilli* (Bhardwaj et al. 2010; Palkovicsné Pézsa et al. 2022) and bolster the host’s immune defenses to decrease infection in the host (Khalkhali and Mojgani 2017). It has been also reported that *E. faecium* can eliminate or inhibit pathogens through various mechanisms, including auto-aggregation with pathogens, adhesion to intestinal mucosa, enhancement of epithelial barrier function, and the production of antimicrobial compounds such as bacteriocins and lysozymes (Klingspor et al. 2015; Qiu et al. 2022). In addition to its pathogen-inhibiting mechanisms, *E. faecium* also contributes to host health by synthesizing essential vitamins that support cellular metabolism, enhance immune response, and promote intestinal homeostasis (Dimidi et al. 2019; Holzapfel et al. 2018).

The advancement of genome sequencing technology has significantly accelerated probiotic research by offering more comprehensive insights and facilitating precise discoveries. High-throughput whole genome sequencing (WGS) technology in probiotics research is essential for identifying genes responsible for beneficial metabolic functions, such as the production of AMPs, vitamins, and enzymes that promote gut health (Hoque et al. 2024a; Hoque et al. 2024b; Rahaman et al. 2024). It also helps to uncover genetic traits related to immune modulation, pathogen inhibition, and antibiotic resistance, ensuring the safety and efficacy of probiotic strains (Hasnat et al. 2024b). This detailed genomic insight supports the development of tailored probiotics with enhanced therapeutic potential for improving human and animal health. Although, several studies have shown that lactic acid bacteria can compete with mastitis-causing pathogens (Kober et al. 2022; Rahman et al. 2024b), the antimicrobial effectiveness of *E. faecium* against bovine mastitis causing pathogens remains unclear. This study therefore sought to isolate and characterize *E. faecium* MBBL3 from the milk of a healthy lactating cow, followed by genome sequencing and comprehensive genomic analysis to assess its probiotic potential and antimicrobial activity against mastitis pathogens.

## Materials and Methods

### Bacterial strain isolation and culture conditions

Milk samples were aseptically collected from healthy lactating cows at the BSMRAU dairy farm (24.09° N, 90.41° E) in Gazipur, Bangladesh. The health status of the cows was confirmed by conducting a California Mastitis Test to ensure the absence of subclinical or clinical mastitis (Hoque et al. 2015). Following collection, the milk samples were enriched in de Man, Rogosa, and Sharpe (MRS) broth (HiMedia, India) and incubated aerobically at 37°C for 24h. The enriched samples were streaked onto MRS agar plates and subjected to further incubation at 37°C for 48 h to facilitate bacterial growth. Colonies obtained from the agar plates were analyzed for species identification using the VITEK 2 system (version 9.01) (Kumaran et al. 2023). Pure colonies from MRS agar were diluted in 0.45% saline to achieve a 0.5 McFarland standard and subsequently inoculated into VITEK 2 system cards. The system provided species-level identification of the isolates within 3 h, confirming the presence of *E. faecium* based on their distinctive biochemical properties, with MBBL3 identified as one of the confirmed isolates. Finally, *E. faecium* MBBL3 isolate was stored in MRS broth at -80 °C supplemented with 20% glycerol (Rahman et al. 2024a).

### Genome sequencing, assembly and annotation of the *E. faecium* MBBL3

The MBBL3 isolate was subcultured overnight at 37°C in nutrient broth (Biolife™, Italy). Genomic DNA extraction was performed using the QIAamp DNA Mini Kit (QIAGEN, Germany). Concentration and purity of the extracted DNA were assessed with a NanoDrop 2000 spectrophotometer (Thermo Fisher Scientific, USA) (García Alegría et al. 2020). Whole-genome libraries were prepared from 1 ng of DNA using the Nextera™ DNA Flex library preparation kit (Illumina, San Diego, USA) and sequenced on an Illumina MiSeq platform (Illumina, USA) using the 2 × 250 bp protocol (Rahman et al. 2024a). The paired-end raw reads (N = 1,261,702) were processed by trimming with Trimmomatic v0.39 (Bolger et al. 2014) and assessed for quality with FastQC v0.11.7 (Andrews 2017). Reads with phred scores greater than 20 (Hoque et al. 2023) were assembled using SPAdes v3.15.6 (Bankevich et al. 2012). Genome annotation was carried out using the NCBI Prokaryotic Genome Annotation Pipeline (PGAP) v6.6 (Tatusova et al. 2016) in combination with the *Enterococcus* CheckM v1.2.2 marker set (Parks et al. 2015).

Subsystem analysis was conducted using the Rapid Annotation using Subsystem Technology (RAST) web server (http://rast.nmpdr.org/) (Overbeek et al. 2014). The core genome multi-locus sequence typing (cgMLST) of the MBBL3 genome was predicted using the BacWGSTdb 2.0 platform, employing a seven-gene MLST scheme targeting *atpA*, *ddI*, *gdh*, *purK*, *gyd*, *pstS*, and *adk* (Feng et al. 2021). Visualization of the MBBL3 genome was performed using Genovi v0.2.16 (Cumsille et al. 2023), while plasmid detection was carried out with PlasmidFinder (Carattoli et al. 2014).

### Evolutionary phylogenetic analysis and ortholog identification of *Enterococcus* strains

The assembled genome of MBBL3 was used to conduct an evolutionary phylogenetic analysis, comparing it with fourteen closely related *Enterococcus* strains (**Table S1**) obtained from the National Center for Biotechnology Information (NCBI) database (https://www.ncbi.nlm.nih.gov/). A maximum likelihood phylogenetic tree was constructed using the Type Strain Genome Server (TYGS; https://tygs.dsmz.de/) (Meier-Kolthoff and Göker 2019), including closely related genomes obtained from the NCBI database. Based on phylogenetic relatedness, five closely related Enterococcus strains namely *E. faecium* MBBL3, *E. faecium* HY07, *E. faecium* BIOPOP-3ALE, *E. faecium* BIOPOP-3WT, and *E. lactis* DH9003 were selected for orthologous gene group prediction using OrthoFinder v2.5.5 (Emms and Kelly 2015), with default parameters. This analysis facilitated the identification of conserved and strain-specific orthogroups, providing a detailed comparison of the genomic content and evolutionary relationships among the *Enterococcus* strains along with MBBL3.

### OrthoANI, pangenome and comparative genome analyses of the *E. faecium* MBBL3

Based on phylogenetic relationships and orthologous gene content, aforementioned five closely related strains were chosen for OrthoANI, pangenome, and comparative genome analyses.

Genomic similarity between *E. faecium* MBBL3 and the other four strains was assessed by calculating Average Nucleotide Identity (ANI) using OrthoANI (Lee et al. 2016) with default parameters. To further investigate genetic relationships, pangenome analysis of the selected strains was conducted using Roary v3.11.2 (Sitto and Battistuzzi 2020), categorizing genes into four distinct classes: ‘core’ (99% ≤ strains ≤ 100%), ‘soft core’ (95% ≤ strains < 99%), ‘shell’ (15% ≤ strains < 95%), and ‘cloud’ (0% ≤ strains < 15%). The Mauve alignment web tool (Darling et al. 2010) was employed to assess synteny across large genomic blocks between *E. faecium* MBBL3 and *E. faecium* HY07. Furthermore, the BLAST Ring Image Generator (BRIG) v 0.95 (Camprubí-Font et al. 2018) was utilized to align and visualize the genomes of five strains, with *E. faecium* MBBL3 (GenBank Accession No. <u>JAZIFO000000000</u>) serving as the reference. Nucleotide alignments were conducted using the BLASTn algorithm, applying an e-value cutoff of 1e-5 to ensure high-confidence matches. The alignment results were visualized through BRIG v0.95, providing a comprehensive view of nucleotide-level similarities and variations across the genomes.

### Analysis of carbohydrate utilization, enzyme profiles, and metabolic pathways in *E. faecium* MBBL3

Multiple biochemical assays (**Table 1**) were conducted in accordance with established protocols (Jung et al. 2012) to characterize the carbohydrate fermentation profiles and enzymatic activities of the *E. faecium* MBBL3 isolate. To further analyze the carbohydrate metabolism of MBBL3, genes associated with carbohydrate-active enzymes (CAZymes) were identified using the Carbohydrate-Active Enzymes (CAZy) database ( http://www.cazy.org/). Protein sequences from the MBBL3 genome were annotated through HMMER v3.4 (Finn et al. 2011) using the dbCAN server (https://bcb.unl.edu/dbCAN2/index.php), with alignments performed against the CAZy database. CAZyme classification was determined based on cut-off thresholds of 1e-15 and 0.35, ensuring high-confidence identification of relevant enzyme families. To predict metabolic pathways and classify genome functions, annotation was performed using the KEGG database (Kanehisa et al. 2016) through the KEGG Automatic Annotation Server (KAAS, https://www.genome.jp/kegg/kaas/) and BlastKOALA (https://www.kegg.jp/blastkoala/).

**Table 1.**
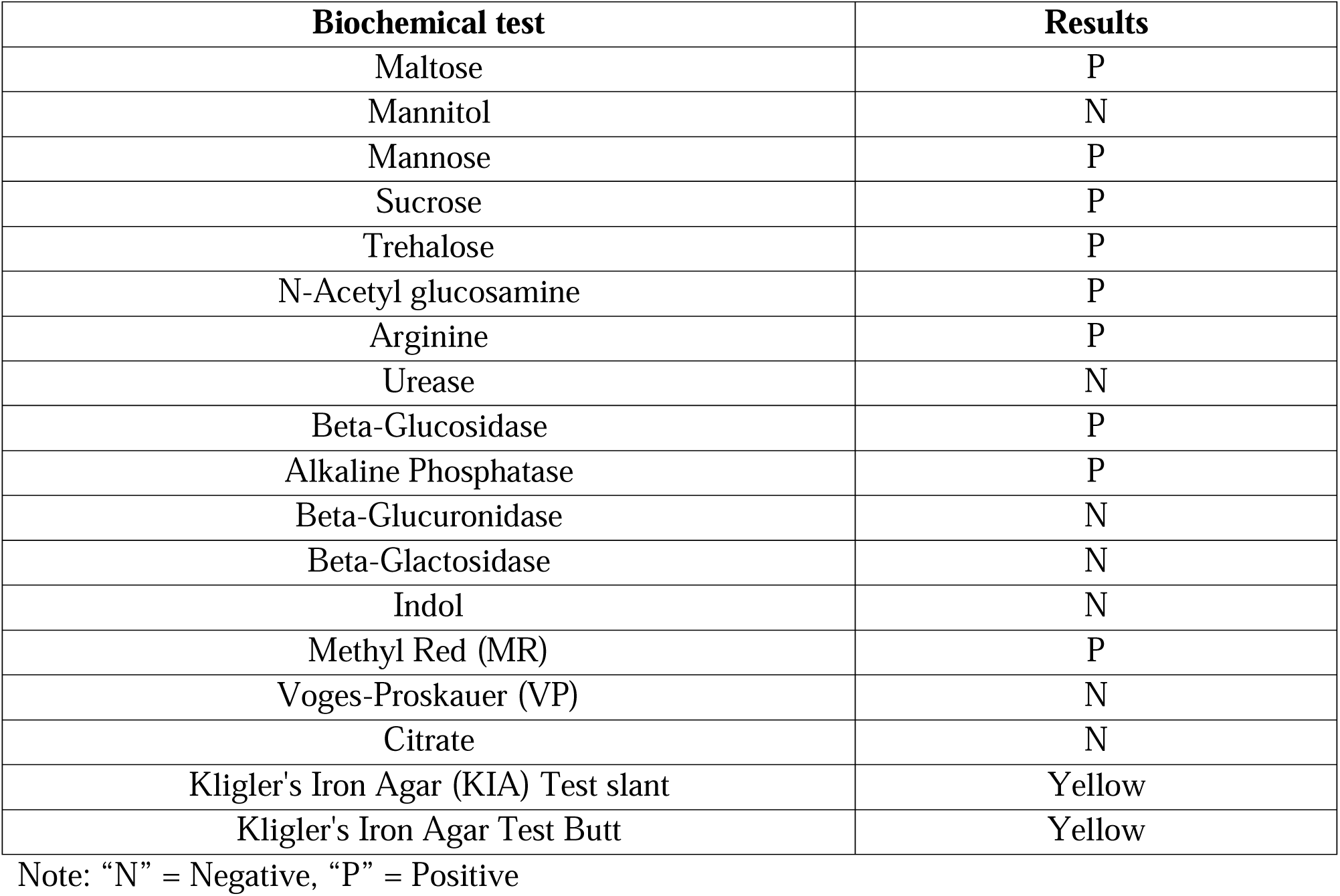
Carbohydrate fermentation activities of the *E. faecium* MBBL3 isolate.

### Analysis of primary and secondary metabolites in *E. faecium* MBBL3

The *E. faecium* MBBL3 genome was further analyzed for primary and secondary metabolite biosynthetic gene clusters using the gutSMASH v1.0 (Pascal Andreu et al. 2021) and antiSMASH v7.0 (Blin et al. 2023) databases, respectively. Bacteriocins and their associated genes were subsequently identified using the BAGEL5 web tool (http://bagel5.molgenrug.nl/index.php; 15 November 2024). To ensure the accuracy of the annotations, the identified bacteriocin domains were manually validated using BLASTp (https://blast.ncbi.nlm.nih.gov/Blast.cgi), confirming high-confidence identification of these functional elements.

### *In-vitro* and genomic safety profiling of *E. faecium* MBBL3

The antibiotic susceptibility of the MBBL3 isolate was assessed using the disc diffusion method (Hoque et al. 2024a). Ten commercially available antibiotics targeting different mechanisms of action were tested, including protein synthesis inhibitors (azithromycin, 15 μg; amikacin, 30 μg; and gentamicin, 10 μg), cell wall synthesis inhibitors (cefepime, 30 μg; ceftriaxone, 30 μg; cefuroxime sodium, 30 μg; imipenem, 10 μg; and meropenem, 10 μg), and DNA gyrase inhibitors (nalidixic acid, 30 μg; and ciprofloxacin, 5 μg) (Oxoid, UK). Briefly, overnight cultures (approximately 10^9^ CFU/mL) were swabbed onto Mueller Hinton agar plates. Antibiotic discs were placed aseptically, and the plates were incubated at 37°C for 24 h. The zone of inhibition (ZOI) was measured, and the susceptibility was classified as resistant (R), intermediate (I), or susceptible (S) following the Clinical and Laboratory Standards Institute (CLSI) M100 guidelines, 2023 (Rai et al. 2023). All tests were performed in triplicate to ensure reproducibility. Antibiotic resistance genes (ARGs) were predicted using the ResFinder 4.0 (Bortolaia et al. 2020) database, while virulence factor genes (VFGs) were identified through the VirulenceFinder 2.0 (Liu et al. 2022) database. Both analyses were conducted using the ABRicate biotool (https://github.com/tseemann/abricate) with default parameters. The hemolytic activity of MBBL3 was assessed by streaking the isolate on blood agar plates supplemented with 5% (v/v) sheep blood. After incubation at 37°C for 24 h, the plates were examined for hemolytic activity, including α, β, or γ hemolysis, with an uninoculated control plate serving as negative control (Oh and Jung 2015). Prophage regions in the MBBL3 genome were identified using the PHASTEST web tool (https://phastest.ca/) (Wishart et al. 2023), and CRISPR loci along with associated CRISPR-associated (Cas) proteins were predicted using CRISPRCasFinder v4.2.20 (Couvin et al. 2018).

### *In-vitro* antimicrobial efficacy of *E. faecium* MBBL3 against mastitis pathogens

To assess the in-vitro antimicrobial efficacy of *E. faecium* MBBL3, concentrated cell-free supernatant (CFS) was prepared according to established protocols with slight modifications (El Oirdi et al. 2021; Zaghloul and Halfawy 2024). MBBL3 was initially cultured in 10 mL nutrient broth (Oxoid, UK) and incubated statically at 37°C for 24 h. A 1 mL aliquot of this overnight culture was transferred to 100 mL nutrient broth and incubated at 37°C for an additional 48 h. The culture was then centrifuged at 6000 rpm for 15 min at 4°C, and the resulting supernatant was neutralized to pH 7.0. The CFS was subsequently filtered through a 0.22 μm Millipore filter to obtain a sterile preparation. The antimicrobial activity of the MBBL3 CFS was assessed against two bovine mastitis pathogens, *Kp* MBBL2 (<u>JBINJS000000000</u>) and *Ec* MBBL4 (<u>JBINJR000000000</u>), using the agar well diffusion assay. Bacterial inocula of *Kp* MBBL2 and *Ec* MBBL4 (10 CFU/mL each) were evenly spread onto nutrient agar plates (Oxoid, UK). Wells (4 mm in diameter) were created in the agar and filled with 60 μL of sterile CFS. A control plate without CFS was maintained for comparison. After incubation at 37°C for 24 h, antimicrobial efficacy was determined by measuring the ZOI surrounding the wells.

### Identification of essential and virulence proteins in mastitis pathogens

Furthermore, *Kp* MBBL2 and *Ec* MBBL4 were used as the target pathogens to investigate the antimicrobial efficacy of *E. faecium* MBBL3 at the molecular level. Essential proteins from these pathogens were identified through BLASTp searches against the Database of Essential Genes (DEG) (Luo et al. 2021) using genome assemblies of *Kp* MBBL2 and *Ec* MBBL4 as queries. BLASTp analyses were conducted with an e-value cutoff of 10 ³ , a sequence identity threshold of ≥50%, and a bit score of ≥100 (Hasnat et al. 2024a). Identified essential proteins were subsequently screened for virulence factors (or genes) using the Virulence Factor Database (VFDB) (Chen et al. 2005) employing the same parameters. The resulting proteins were considered essential for both bacterial survival and pathogenicity (Hasnat et al. 2024b).

### Molecular interactions between Enterolysin_A and virulence proteins of mastitis pathogens

The antimicrobial efficacy of Enterolysin_A was assessed by targeting essential and virulence proteins of *Kp* MBBL2 and *Ec* MBBL4, resulting in the identification of 34 and 32 key proteins, respectively (**Table S2**). Protein structures for both pathogens were retrieved from AlphaFold and preprocessed using Discovery Studio 2024 (Jejurikar and Rohane 2021). Similarly, the structure of Enterolysin_A was obtained from AlphaFold and prepared for analysis. Molecular docking studies were performed using the Maestro BioLuminate (v.5.5) software package from Schrödinger (Islam et al. 2024) to evaluate interactions between Enterolysin_A and the identified proteins. Remarkably, Enterolysin_A demonstrated a strong binding affinity to the outer membrane usher protein, type 1 fimbrial synthesis in mastitis pathogenesis (Gato et al. 2020; Olson et al. 2024). To explore the biological implications of this interaction, a membrane system was constructed. The Enterolysin_A-usher protein complex was embedded within this system, and molecular dynamics simulations (MDS) were performed over 200 ns to investigate the stability and behavior of the interaction under dynamic conditions (Hasnat et al. 2024b).

## Results

### Genomic features of the *E. faecium* MBBL3

The draft genome of *E. faecium* MBBL3 spans 2,971,195 base pairs (bp) across 106 contigs, with a genome coverage of 95.61x and a GC content of 37.97%, indicating robust sequencing coverage and balanced nucleotide composition. It includes two plasmid replicons harboring repUS15 (1,041 bp) and rep1 (1,417 bp). CheckM analysis revealed the MBBL3 genome to be 98.02% complete with minimal contamination (0.17%). Based on cgMLST, the *E. faecium* MBBL3 genome belonged to *Enterococcus* sequence type 224 (ST224). **Table 2** provides detailed genomic features of *E. faecium* MBBL3, including coding sequences (CDs), RNAs, CRISPR regions, Cas clusters, prophages, ARGs, VFGs, and number of subsystems. The circular genome representation of MBBL3 (**Fig. 1A**), along with the classification of its genes into clusters of orthologous genes (COGs) (**Fig. 1B**), underscores the genomic framework of MBBL3, revealing key genes involved in fundamental biological processes critical for its growth, survival, and functional capabilities.

**Fig 1.**
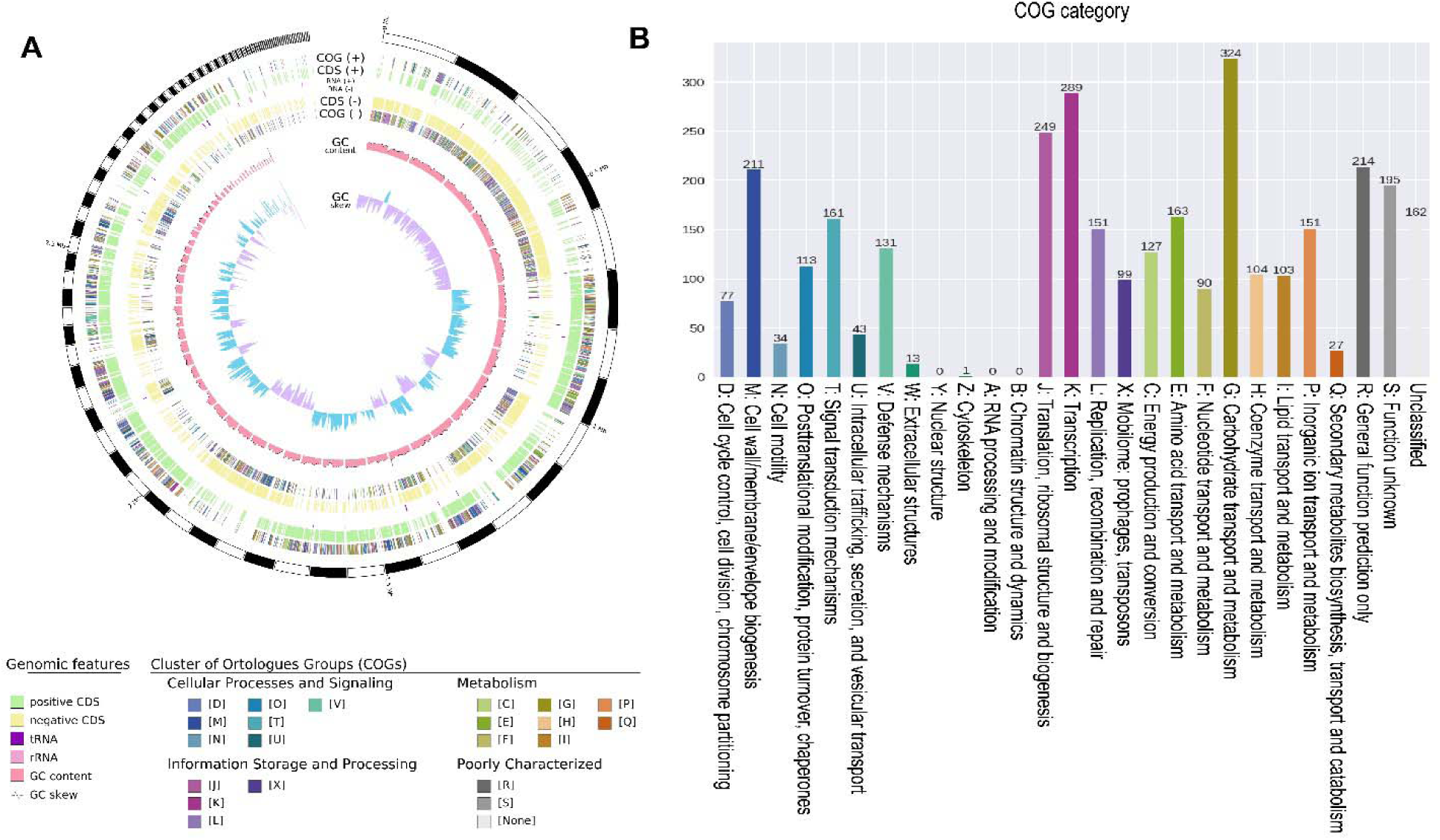
Genomic architecture and functional categorization of *E. faecium* MBBL3. (A) Circular genome map of *E. faecium* MBBL3 visualized from the outside to the center: genome contigs, coding sequences (CDS) on forward strand with clusters of orthologous groups (COG) category annotation, tRNA and rRNA, CDS on reverse strand with COG category annotation, GC content, and GC skew. (B) Functional distribution of CDS based on COG categories predicted in *E. faecium* MBBL3 genome.

**Table 2.** Key genomic features of *E. faecium* MBBL3.

| Features | Genome information |
| --- | --- |
| Genome size (bp) | 2,971,195 |
| No. of contigs | 106 |
| Genome coverage (x) | 95.61 |
| GC content (%) | 37.97 |
| Genome completeness (%) | 98.02 (0.17% contamination) |
| Sequence type (ST) | 224 |
| *N*<sub>50</sub> value (bp) | 72,781 |
| Total genes | 3,030 |
| Protein coding genes | 2,843 |
| Total coding sequences | 2,954 |
| tRNA | 63 |
| rRNA | 07 |
| CRISPR | 03 |
| Cas proteins | 09 |
| Prophages | 0 |
| Number of plasmids | 03 |
| Antibiotic resistance gene | 02 |
| Virulence genes | 01 |
| Pseudo gene | 111 |

### Evolutionary analysis of MBBL3 shows high genetic resemblance to other *E. faecium* strains

The evolutionary phylogenetic analysis of *E. faecium* MBBL3, conducted in comparison with 14 other *Enterococcus* strains (**Table S1**), demonstrated a close genetic relationship with strains from diverse environments. Notably, MBBL3 exhibited high similarity to *E. faecium* HY07, previously isolated from Chinese sausages (Duan et al. 2019), *E. faecium* BIOPOP-3WT and BIOPOP-3ALE from fermented dairy products (Min et al. 2020), and *E. lactis* DH9003 from goat milk (Yang et al. 2023) (**Fig. 2A**). Homologous gene content analysis of these closely related genomes revealed 12,715 genes, of which 97.2% were classified into 2,738 orthogroups. Among these, 1,990 orthogroups were classified as core orthogroups, representing highly conserved genetic elements shared across the strains. In contrast, 1,825 orthogroups were categorized as accessory orthogroups, reflecting genomic plasticity and strain-specific variation. Notably, MBBL3 was present in 2,492 orthogroups (91% of total), with 11 orthogroups identified as species-specific, suggesting unique genetic features of this strain. OrthoANI analysis revealed that MMBL3 genome shared > 98% similarity with all of the reference genomes supporting the results of phylogenetic analysis (**Fig. 2B**). The evolutionary phylogenetic analysis was further supported by pangenome analysis of five closely related strains namely *E. faecium* MBBL3, HY07, BIOPOP-3WT, BIOPOP-3ALE, and *E. lactis* DH9003. The pangenome matrix, based on gene presence and absence, revealed clusters of genes and a dendrogram showing their relationships (**Fig. 3A**). A total of 4,260 genes were predicted, including 2,015 core genes present in over 99% of the genomes, highlighting their conservation across strains. Additionally, 2,245 shell genes, found in 15% to 95% of the genomes, showed moderate variability. No cloud genes (present in < 15% of the genomes) or soft-core genes were detected, indicating a relatively stable genomic composition (**Fig. 3B**).

**Fig 2.**
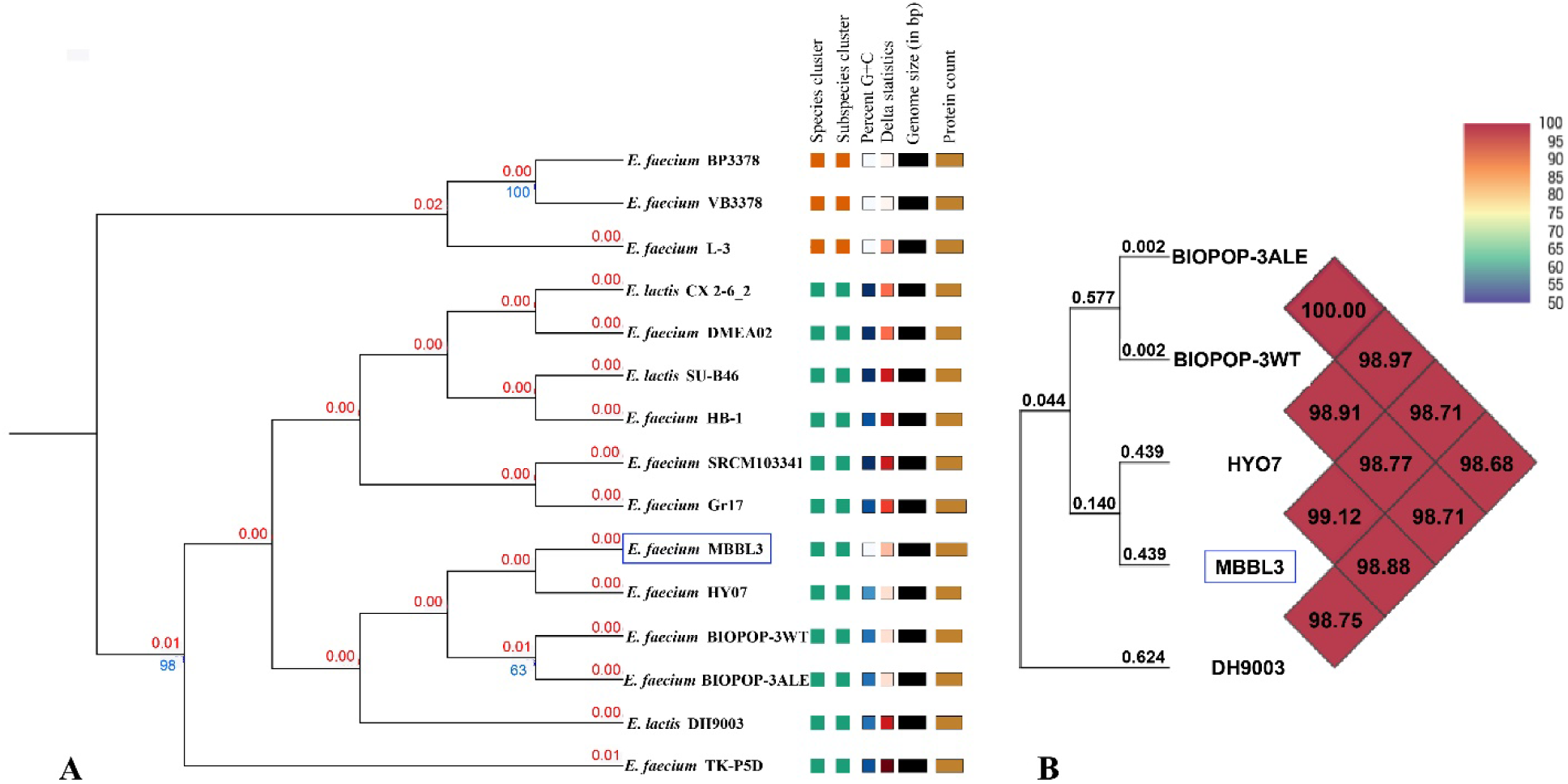
Phylogenomic classification of *E. faecium* MBBL3 based on: (A) Genome Basic Local Alignment Search Tool (BLAST) distance phylogeny (GBDP) using Type Strain Genome Server (TYGS) platform. Tree was inferred with FastME 2.1.6.1 (94) from GBDP distances calculated from genome sequences. The branch lengths are scaled in terms of GBDP distance formula D5. The numbers above branches are GBDP pseudo-bootstrap support values > 60 % from 100 replications, with an average branch support of 9.8 %. The tree was rooted at the midpoint (95). (B) OrthoANI values using Orthologous Average Nucleotide Identity Tool (OAT) software (https://www.ezbiocloud.net/tools/orthoani). Heatmap presents OrthoANI values of *E. faecium* MBBL3 and four closely related strains of *E. faecium* and *E. lactis*. The genome of the *E. faecium* MBBL3 (JAZIFO000000000.1) is highlighted on blue box.

**Fig 3.**
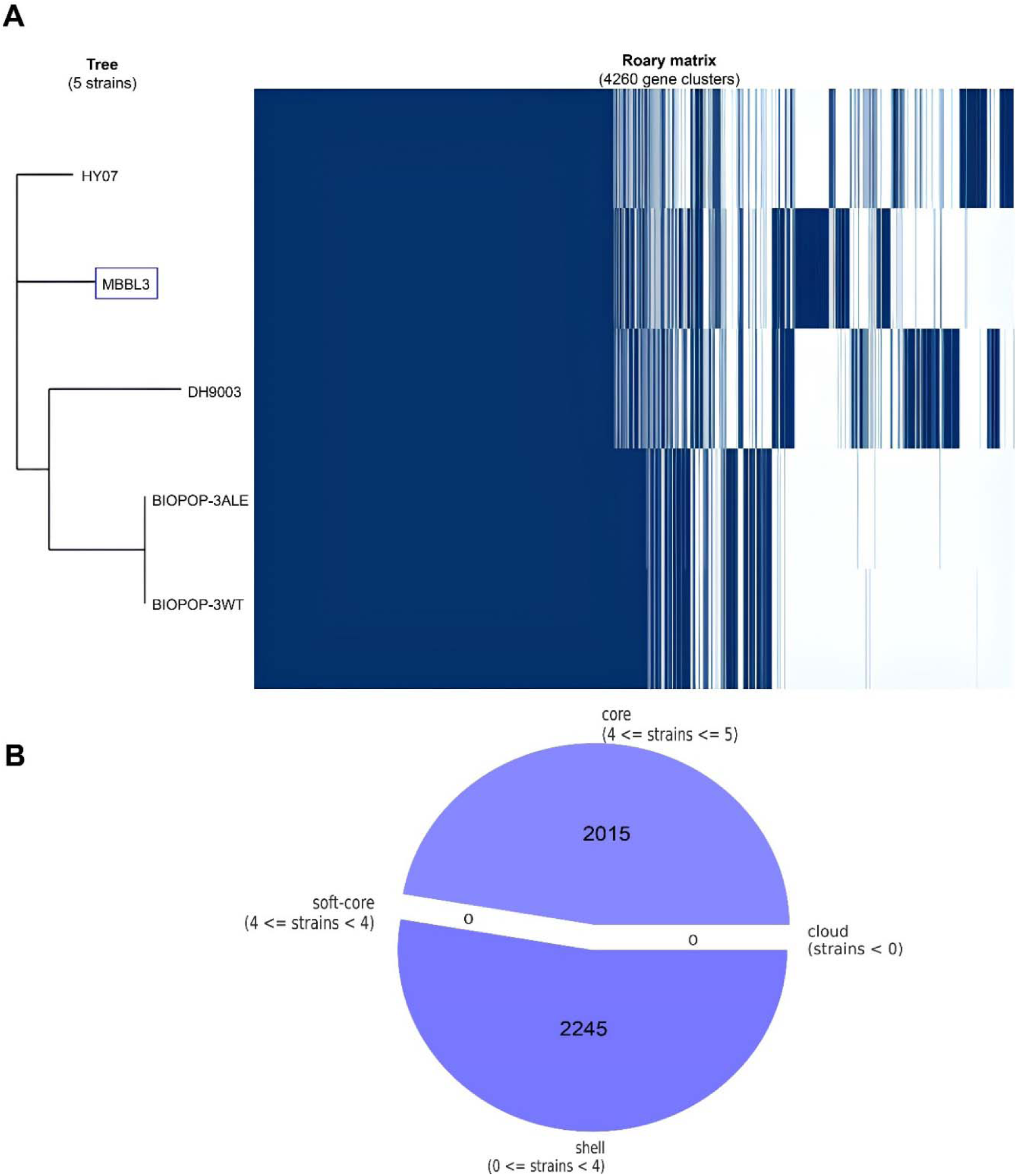
Pangenome analysis of *E. faecium* strains. (A) Pangenome based (gene presence and absence) gene clustering matrix of *E. faecium* MBBL3 (enclosed in red box) and four closely related genomes of *E. faecium* and *E. lactis* from Bangladesh and beyond. (B) Breakdown of genes in five *E. faecium* genomes. We generated the figures based on the data obtained from Roary pangenome analysis using the roary_plots.py script and genes available in the five genomes of *E. faecium*.

### *E. faecium* MBBL3 exhibits minimal genomic discrepancy with reference genomes

The *E. faecium* strain HY07, identified as the closest relative to *E. faecium* MBBL3, was selected for multiple genome alignment to assess genomic synteny. The analysis revealed a high degree of synteny between the genomes of MBBL3 and HY07 (**Figure 4A**). Furthermore, comparative analysis using the BRIG tool demonstrated substantial conservation of genomic regions, visualized as solid circular segments in varying colors, while areas of variability were indicated as blank segments (**Fig. 4B**). The genomic maps revealed minimal large-scale structural variation among the strains, with homologous regions surrounding the reference genome exhibiting more than 95% sequence identity.

**Fig 4.**
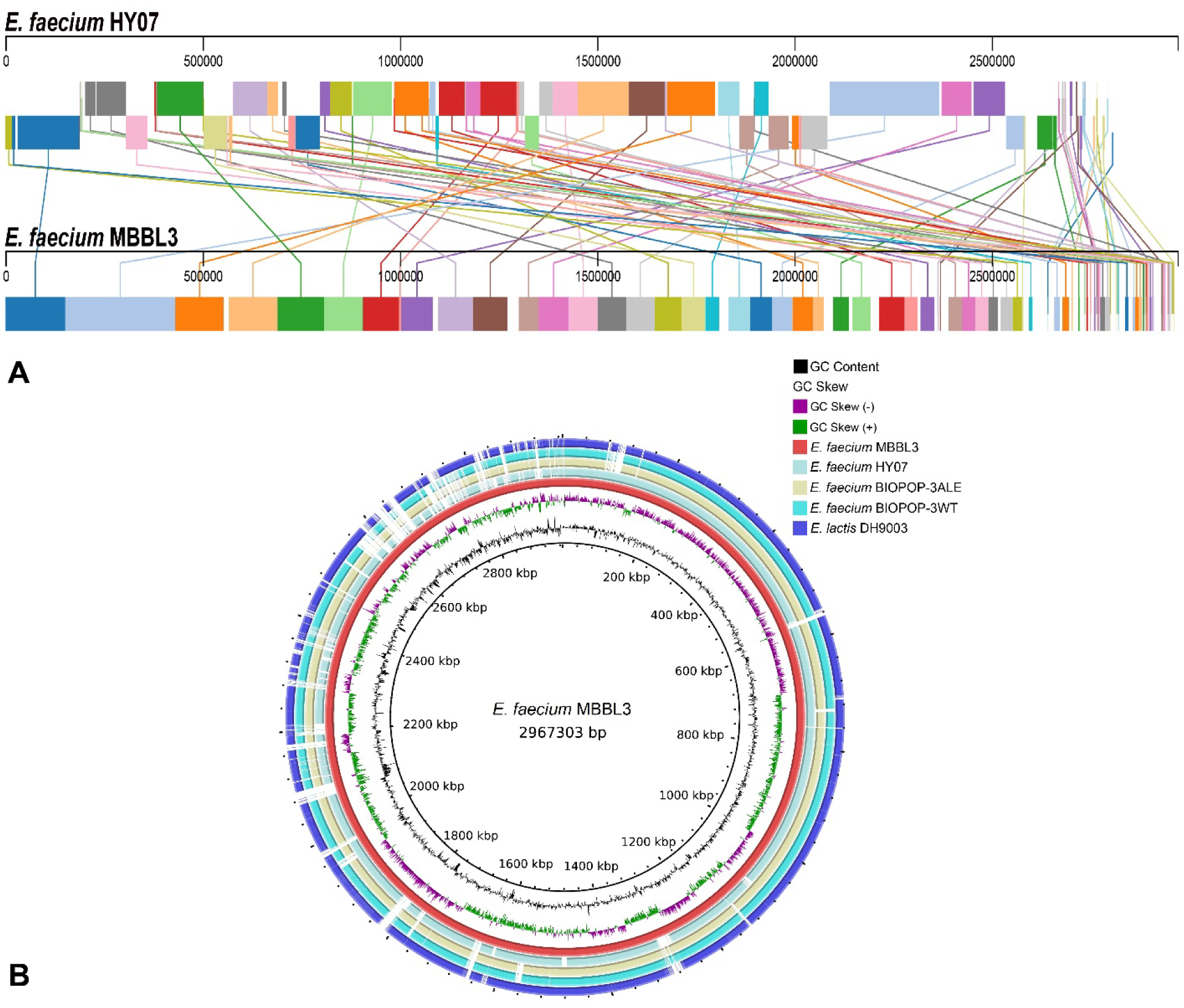
Overview of multiple genome alignment of *E. faecium* strains. (A) Multiple genome alignment of *E. faecium* MBBL3 and *E. faecium* HY07. Each genome sequence is displayed as a horizontal panel with its name labeled on the left. Homologous blocks are represented by the same color and connected within the genome by lines. (B) Nucleotide alignments of five *E. faecium* genomes, generated with the BLASTn with an e-value cut-off 664 1e^-5^ using BRIG 0.95. Circles (from inside to outside) 1 and 2 (GC content; black line and GC skew; purple and green lines), circle 3 (reference *E. faecium* MBBL3 genome; red color), circle 4 (mapped *E. faecium* HY07 genome; cyan color), circle 5 (mapped *E. faecium* BIOPOP-3ALE genome; light brown color), circle 6 (mapped *E. faecium* BIOPOP-3WT genome; sky blue color), and circle 7 (mapped *E. lactis* DH9003 genome; dark blue color).

### *E. faecium* MBBL3 demonstrates capabilities for carbohydrate utilization and enzymatic activities

The *E. faecium* MBBL3 strain demonstrated the capacity to ferment a variety of carbohydrates, including maltose, mannose, sucrose, trehalose, N-acetylglucosamine, and arginine (**Table 1**). Enzymatic assays revealed positive activities for sucrose production, arginine decarboxylation, and β-galactosidase. In contrast, the strain was negative for mannitol fermentation, urease activity, β-glucuronidase, and β-glactosidase activity. The Kligler Iron Agar (KIA) test resulted in uniform yellow coloration in both the slant and butt, indicative of glucose and galactose metabolism. Additionally, the strain did not produce indole, exhibited a negative Voges-Proskauer (VP) reaction, and failed to utilize citrate (**Table 1**). The MBBL3 genome was further analyzed against the CAZy database, revealing a total of 76 genes across six CAZyme classes such as glycoside hydrolases (GH), glycosyltransferases (GT), carbohydrate esterases (CE), carbohydrate binding modules (CBM), auxiliary activities (AA), and polysaccharide lyases (PL) (**Fig. 5**). Among these, the GH (46 genes) family was the most abundant, followed by the GT (20 genes), CE (13 genes), CBM (06 genes), AA (02 genes) and PL (02 genes) family. Importantly, five CAZy families such as GH73, GH18, CBM50, CE4, and auxiliary activity 10 (AA10) were identified for their potential antimicrobial properties, underscoring their functional significance within the MBBL3 genome.

**Fig 5.**
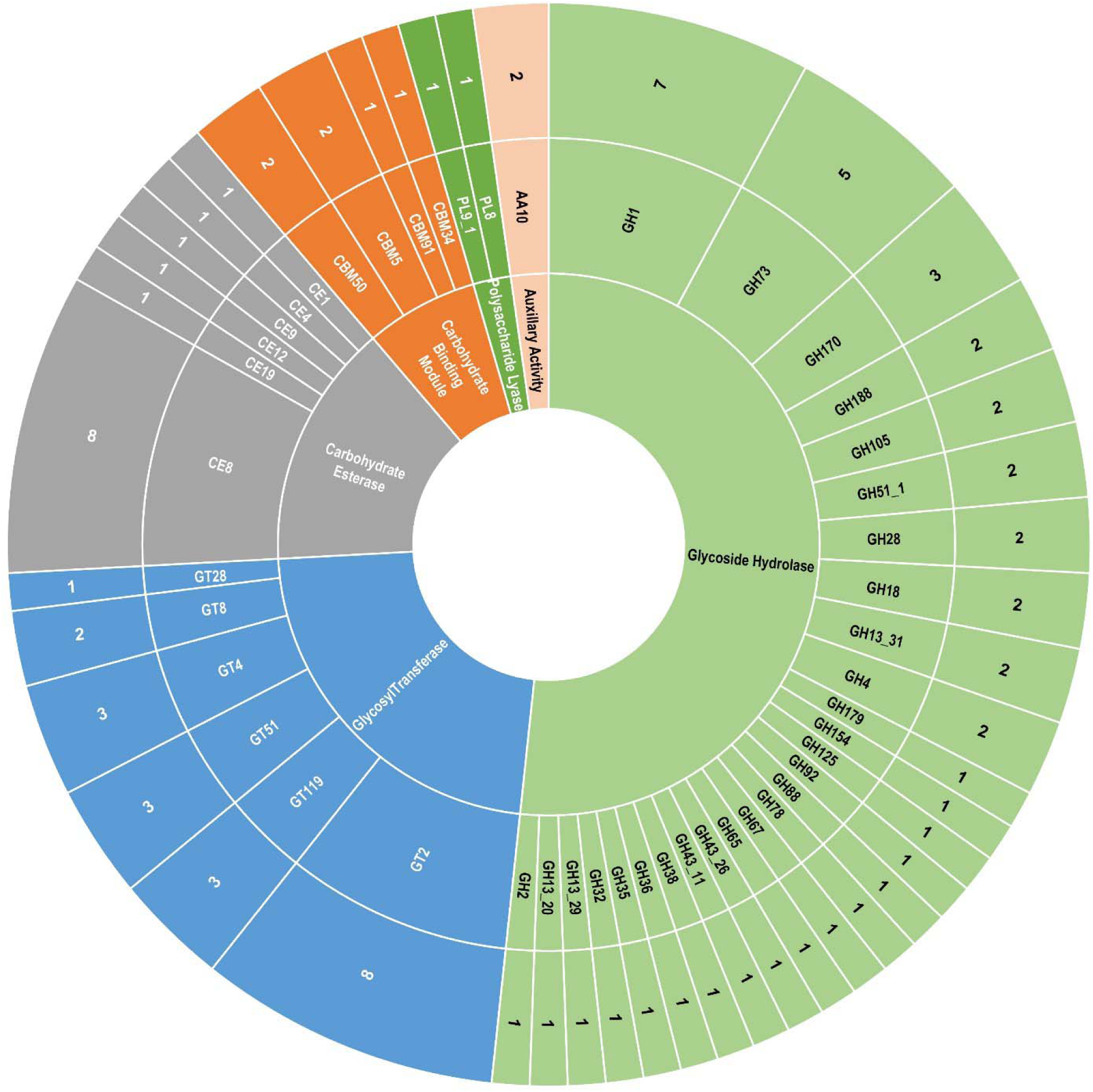
Distribution of CAZymes in the genome of *E. faecium* MBBL3. CAZymes were identified by querying the CAZy database using the dbCAN webserver. Each CAZyme class is represented by distinct colors: pink for auxiliary activity families, green for polysaccharide lyase families, coral for carbohydrate-binding module families, grey for carbohydrate esterase families blue for glycosyl transferase families, and pastel-green for glycoside hydrolase families. The diagram features rings that denote (from the innermost to the outermost) CAZyme classes, CAZyme families, and the number of genes identified within each family, respectively.

### *E. faecium* MBBL3 utilizes different metabolic pathways to facilitate adaptation and survival

To better understand how *E. faecium* MBBL3 metabolizes nutrients, interacts with its environment, and potentially influences health or disease mechanisms, KEGG pathway analysis was conducted. This analysis annotated 1,468 genes, categorized into 39 subcategories across six primary functional groups. The majority of these genes were associated with metabolism (54.29%), followed by genetic information processing (17.22%), environmental information processing (14.70%), human diseases (6.61%), cellular processes (4.16%), and organismal systems (3.02%) (**Fig. 6A**). The primary carbohydrate metabolic pathways in MBBL3 included pyruvate, amino and nucleotide sugar, galactose, fructose, and mannose metabolism, along with glycolysis/gluconeogenesis (**Fig. 6B**). In addition, most abundant amino acid metabolic pathways in MBBL3 were those involving alanine, aspartate, glutamate, cysteine, and methionine (**Fig. 6C**). Further analysis of central carbohydrate metabolism revealed a complete set of 10 genes encoding the Embden-Meyerhof pathway for glycolysis (M00001), enabling the conversion of glucose (α-D-glucose and β-D-glucose) to pyruvate. Additionally, all six core glycolytic genes were also identified in the gluconeogenesis pathway. The genome encoded the pyruvate oxidation module (5 genes, M00307), which catalyzes the oxidative decarboxylation of pyruvate to acetyl-CoA through the pyruvate dehydrogenase complex under aerobic conditions or pyruvate:ferredoxin oxidoreductase under anaerobic conditions. The oxidative phase of the pentose phosphate pathway (M00006) was also present, facilitating the conversion of glucose-6-phosphate to ribulose-5-phosphate. Furthermore, the genome contained genes for phosphoribosyl pyrophosphate (PRPP) biosynthesis (M00005), converting ribose-5-phosphate to PRPP (**Table S3**). As an anaerobic bacterium, MBBL3 encodes L-lactate dehydrogenase (K00016), which catalyzes the reduction of pyruvate to L-lactate under limited oxygen conditions, playing a vital role in anaerobic glucose metabolism:

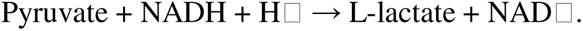

**Fig 6.**
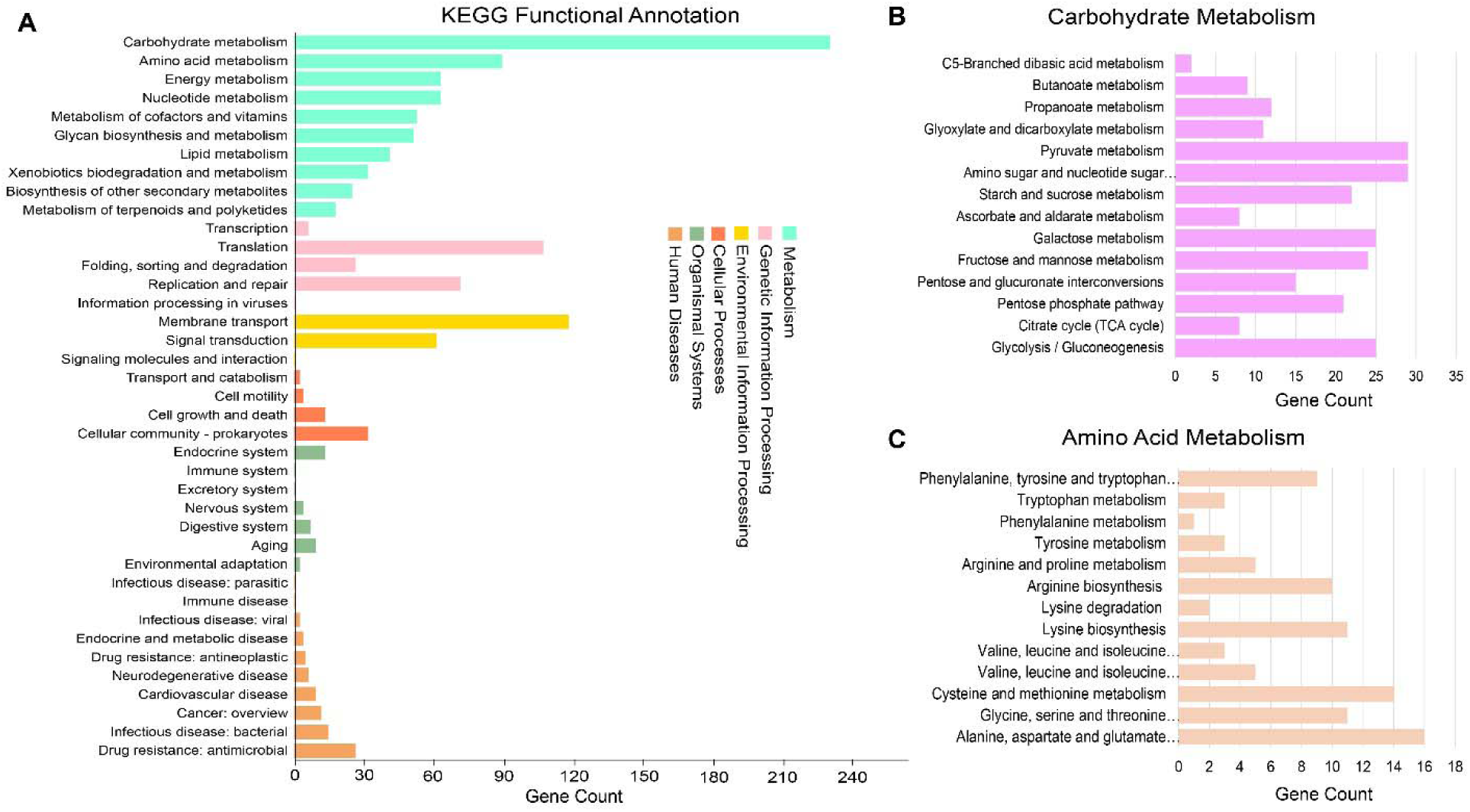
Metabolic functional potentials of *E. faecium* MBBL3 annotated via the KEGG database. (A) Comprehensive overview of functional classes categorized into six main functional categories, each represented by distinct colors: aquamarine for metabolism, pink for genetic information processing, golden for environmental information processing, coral for cellular processes, dark sea-green for organismal systems, and sandy brown for human diseases. (B) Subcategories of carbohydrate metabolism are represented by mauve color horizontal bar chart, and (C) subcategories of amino acid metabolism are represented by double Spanish white color horizontal bar chart.

The genome also encodes anaerobic ribonucleoside-triphosphate reductase (K21636), an oxygen-sensitive enzyme responsible for reducing CTP to dCTP, reflecting adaptation to the oxygen-deprived environment of the human gut. Additionally, genes encoding bile salt hydrolase proteins (K01442) enhance acid and bile salt tolerance, supporting the survival of MBBL3 in the gut. Surface adhesion-related genes (*ltaS*, *tuf*, *yidC*, *gapA*) were also predicted, which facilitate adherence to the intestinal mucosa, promoting colonization stability and contributing to probiotic functionality. The genome further encodes the complete F0F1 ATP synthase subunits (A, B, C, alpha, beta, gamma, delta, and epsilon) and components of the cellobiose PTS system (EIIA, EIIB, and EIIC), which play key roles in acid tolerance and survival under acidic conditions. Moreover, genes linked to heat stress resistance, including *clpB*, *clpX*, *clpP*, *recA*, *HSP20*, *groEL*, and *dnaK*, were identified in MBBL3 genome. The above-described genes, their protein ID, and KO ID are given in **Table 3**. Moreover, six cold-shock proteins (WP_104760464.1, WP_002292860.1, WP_002293303.1, WP_002294067.1, WP_002294871.1, WP_002309330.1) annotated in MBBl3 genome underscores its ability to adapt to environmental stressors, including cold stress.

**Table 3.**
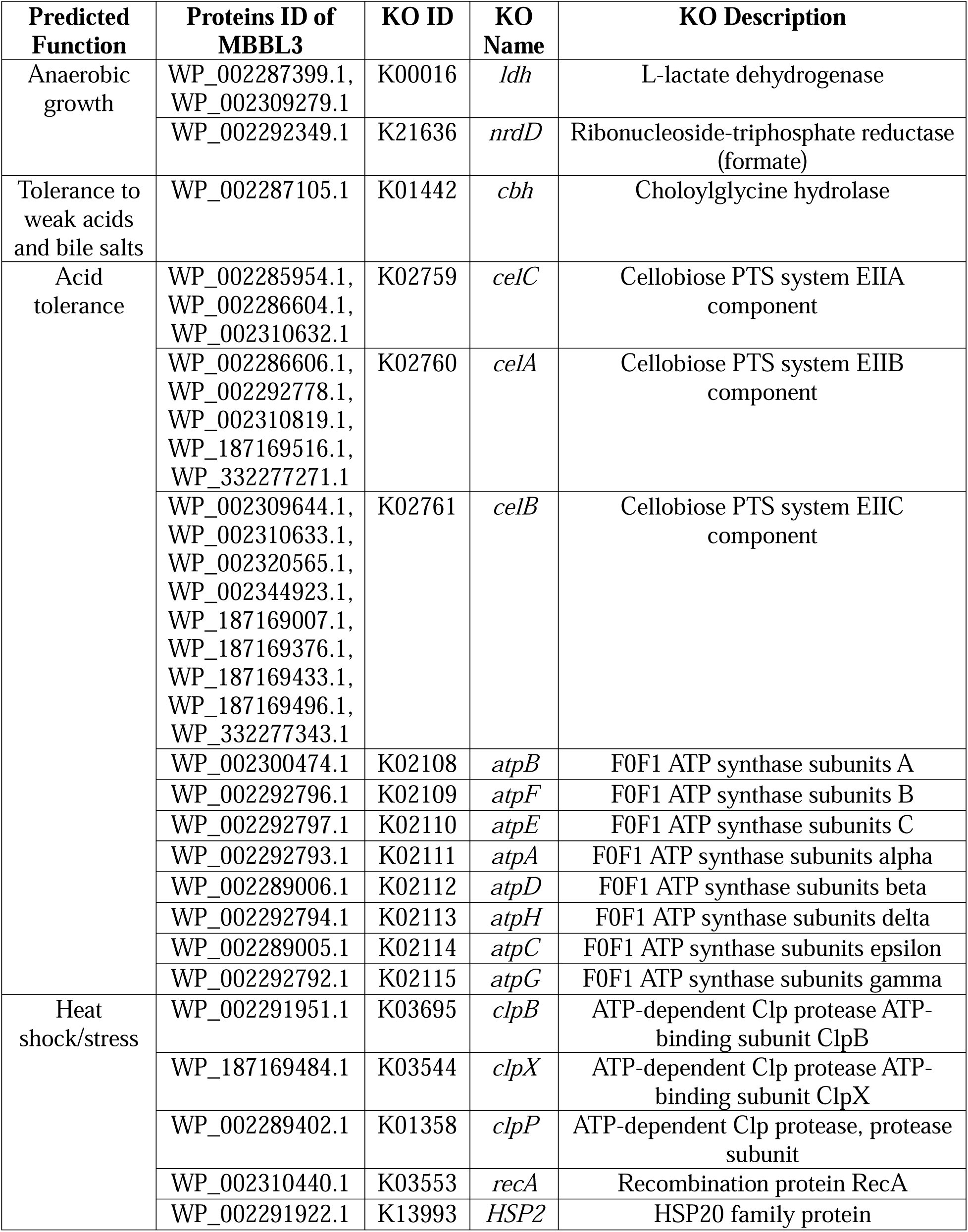
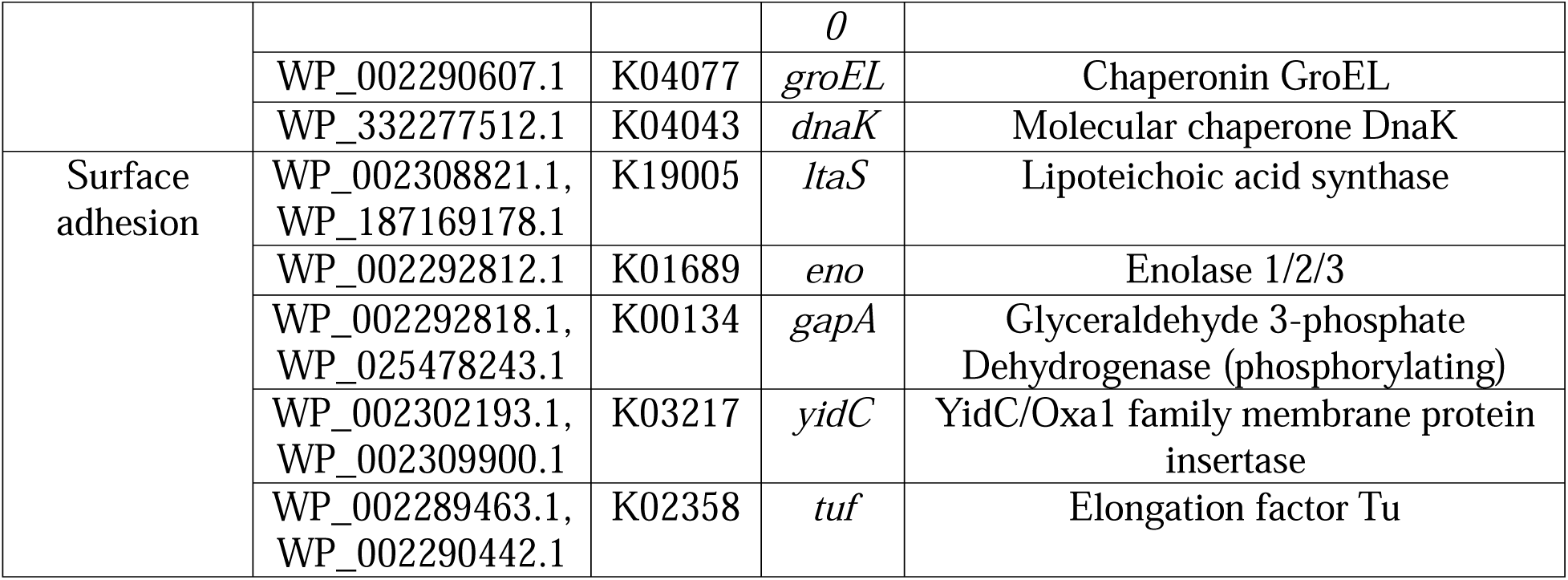
Important probiotic related genes predicted in *E. faecium* MBBL3 genome.

### *E. faecium* MBBL3 possesses gene repertoire for key primary and secondary metabolites

In *E. faecium* MBBL3 genome, we identified four primary (**Figs. 7A-D, Table S4**) and four secondary (**Figs. 7E-H, Table S4**) metabolite regions. The primary metabolic gene clusters identified in the MBBL3 genome includes pyruvate to acetate formate, PFOR II pathway, gallic acid (GA) metabolism (gallic_acid_met) and arginine to hydrogen carbonate (Arginine2Hcarbonate) (**Figs. 7A-D**). The secondary metabolite regions of MBBL3 included Type III polyketide synthase (T3PKS) with a similarity score of 0.27 to 2,4- diacetylphloroglucinol (2,4-DAPG), a type of polyketide (region 1). Rest of the three regions revealed the highest similarity to aborycin (RiPP type), enterocin NKR-5-3B (RiPP type), and sodorifen (terpene type), with similarity scores of 0.20, 0.18, and 0.26, respectively (**Figs. 7E-H** and **Table S5**). Genome analysis for bacteriocin biosynthetic gene clusters (BBGC) identified three BBGC regions in the MBBL3 genome *viz*. contigs 4, 52 and 54 (**Fig. 8**). A sactipeptides class was identified in contig 4, which included key genes such as *LanD*, *BmbF*, and five ABC transporter protein (**Fig. 8A**). Besides, contig 52 contained the biosynthetic gene cluster for the bacteriocin *Enterolysin_A* (**Fig. 8B**), while contig 54 encoded *UviB* (**Fig. 8C**), a core peptide known to be associated with bacteriocin production.

**Fig. 7.**
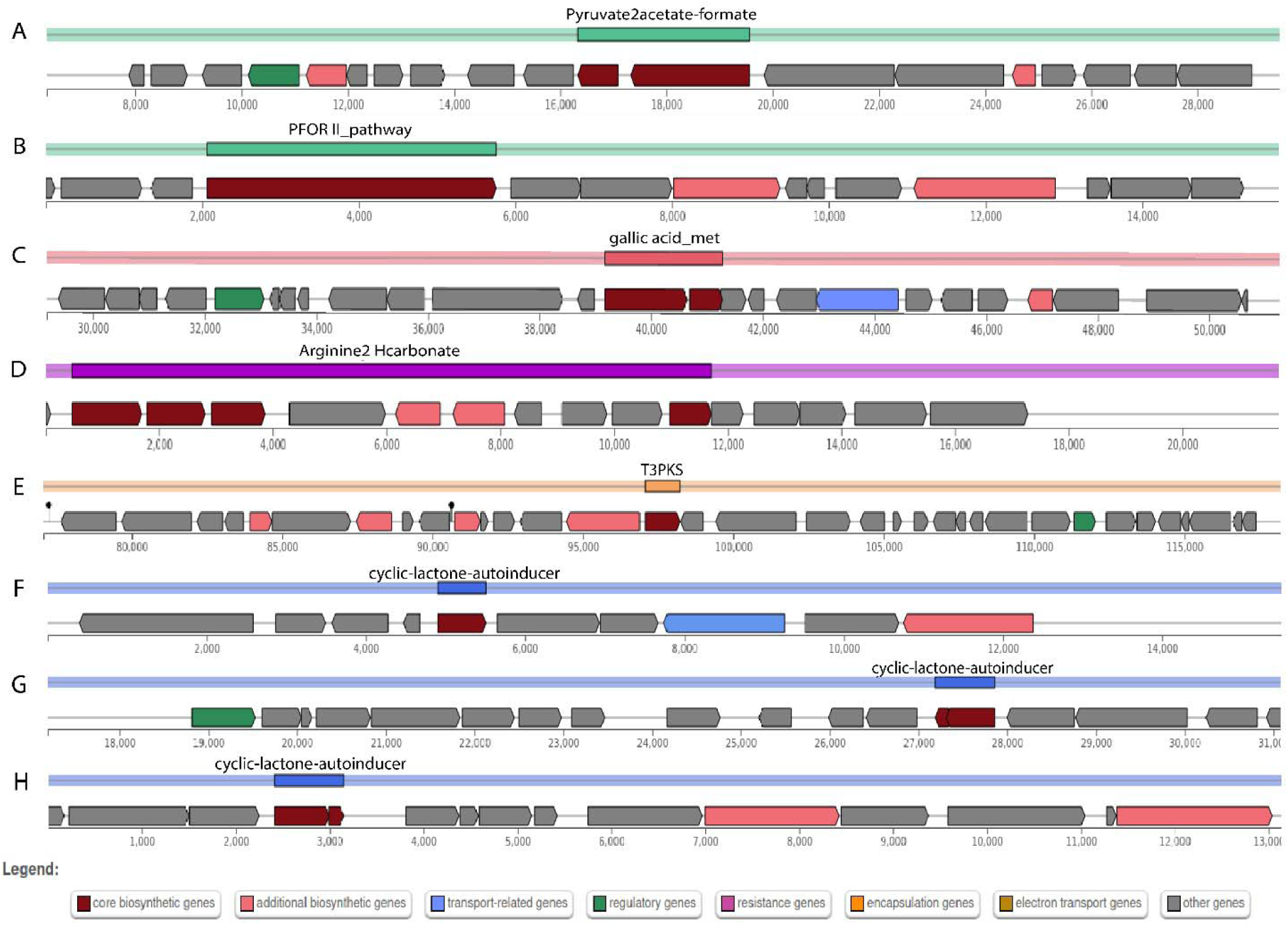
Overview of primary (A-D) and secondary (E-H) metabolite regions in *E. faecium* MBBL3. (A) Pyruvate to acetate formate pathway, (B) PFOR II pathway, (C) Gallic acid metabolism (gallic_acid_met), (D) Arginine to hydrogen carbonate (Arginine2Hcarbonate) metabolism, (E) Type III Polyketide Synthases (T3PKS), and (F), (G), and (H) associated with cyclic-lactone autoinducer pathways.

**Fig. 8.**
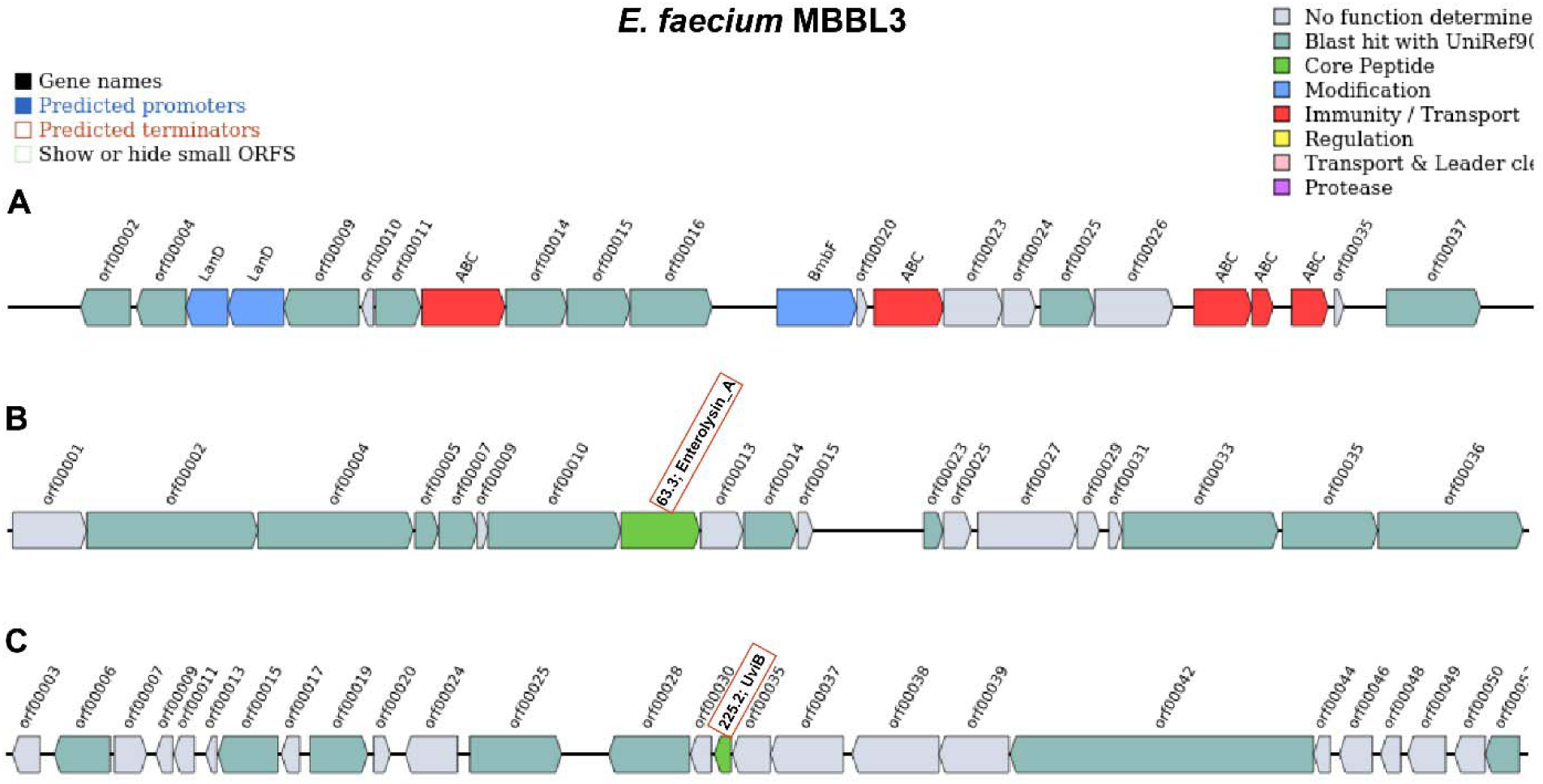
Overview of bacteriocin biosynthetic gene cassettes (BBGCs) in the *E. faecium* MBBL3 genome. (A) Sactipeptides, (B) Enterolysin_A and (C) UviB bacteriocin gene cluster, highlighting their potential roles in antimicrobial therapy and food safety.

### *In-vitro* and genomic analyses suggest the safe use of *E. faecium* MBBL3 as a probiotic

Antibiotic susceptibility testing (AST) revealed that the MBBL3 isolate was susceptible to 50% of the tested antibiotics (5 out of 10), including ceftriaxone, cefuroxime sodium, ciprofloxacin, imipenem, and meropenem. However, the strain demonstrated resistance to azithromycin, amikacin, and gentamicin, intermediate resistance to nalidixic acid, and a susceptible-dose-dependent (SDD) response to cefepime (**Table S6**). Genome annotation identified two ARGs, *aac(6’)-Ii* and *msr(C)*, conferring resistance to aminoglycosides and macrolides, consistent with the AST findings (**Table S7**). Additionally, a single VFG, *acm*, was detected, suggesting low pathogenic potential for MBBL3. Furthermore, MBBL3 exhibited gamma (γ) hemolysis on blood agar, suggesting an inability to lyse red blood cells (**Fig. 9A**). Analysis of mobile genetic elements (MGEs) revealed that the MBBL3 genome lacked prophage regions but contained three CRISPR regions and two Cas clusters (**Table S8**).

**Fig. 9.**
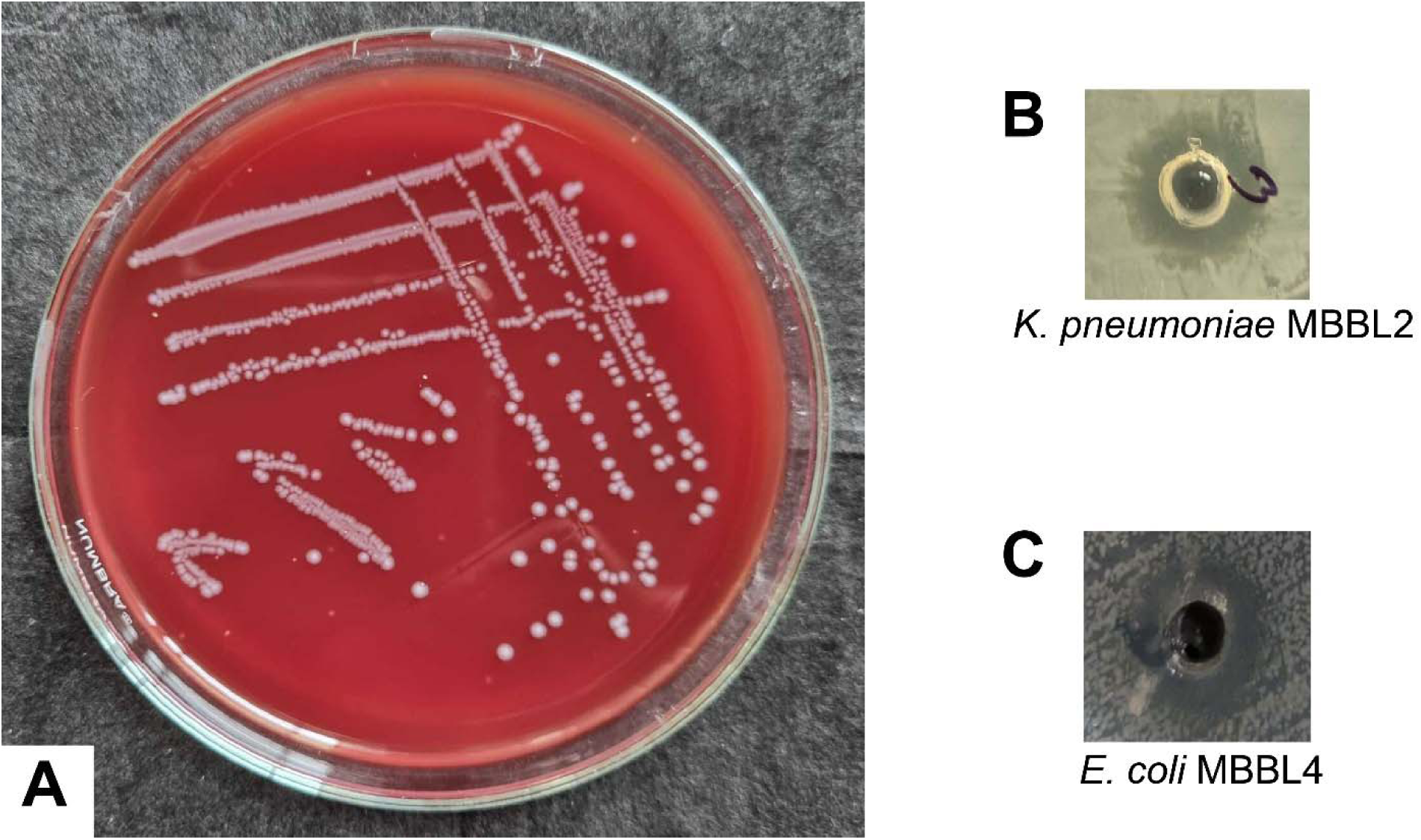
Antimicrobial efficacy of *E. faecium* MBBL3 against mastitis pathogens. (A) Gamma hemolysis of *E. faecium* MBBL3 on the blood agar plate supplemented with 5% (v/v) sheep blood after 24 h of incubation at 37 °C. Antimicrobial effects of cell free supernatant of *E. faecium* MBBL3 against mastitis pathogens (B) *K. pneumoniae* MBBL2 and (C) *E. coli* MBBL4, isolated from milk of clinical mastitis affected cows.

### *E. faecium* MBBL3 exhibits potential antimicrobial efficacy against mastitis pathogens

The *E. faecium* MBBL3 showed effective antimicrobial activity against two major pathogens of bovine mastitis *viz*. *Kp* MBBL2 and *Ec* MBBL4 (Hoque et al. 2024b; Hoque et al. 2020). The *in-vitro* agar well diffusion assay using *E. faecium* MBBL3 revealed inhibition zones of 13.8 mm for *Kp* MBBL2 (**Fig. 9B**) and 14.6 mm for *Ec* MBBL4 (**Fig. 9C**).

### Genome-wide analysis identifies key essential and virulence proteins in mastitis pathogens

Mastitis pathogens, *Kp* MBBL2 and *Ec* MBBL4, were further analyzed to identify essential and virulence-associated proteins. Genome-wide protein analysis revealed 5,131 proteins in *Kp* MBBL2 and 4,196 proteins in *Ec* MBBL4. Screening these proteomes against the DEG identified 209 essential proteins in *Kp* MBBL2 and 289 in *Ec* MBBL4, which were prioritized based on their critical roles in bacterial viability. Further analysis against the VFDB identified 34 virulence-associated proteins in *Kp* MBBL2 and 32 in *Ec* MBBL4 (**Data S1**), highlighting their involvement in the pathogenicity of mastitis in lactating mammals. The identification of proteins with both essential and virulence-associated properties underscore their potential as strategic targets for developing novel antimicrobial therapies.

### Molecular screening reveals interaction between Enterolysin_A of *E. faecium* MBBL3 and usher protein of mastitis pathogens

Enterolysin_A was screened against 34 proteins from *Kp* MBBL2 and 32 proteins from *Ec* MBBL4. Remarkably, Enterolysin_A exhibited strong molecular interactions with the outer membrane usher protein, a critical component of pili biogenesis in both bacterial strains. Detailed analysis revealed that amino acids TYR43, ARG123, ARG125, GLY126, and ILE149 of Enterolysin_A interacted with VAL99, LYS102, PHE131, THR134, ARG147, GLU153, TYR201, GLU208, ASP209, and THR214 of the usher protein. The binding site and orientation of Enterolysin_A suggested that this bacteriocin can obstruct the channel of usher protein, potentially impairing its function (**Fig. 10A**). To assess the structural and functional stability of the Enterolysin_A-usher protein complex, dynamics simulations were performed in a membrane environment. The complex displayed energetically favorable interactions and remained stable throughout 200 ns of MDS (**Fig. 10B**).

**Fig. 10.**
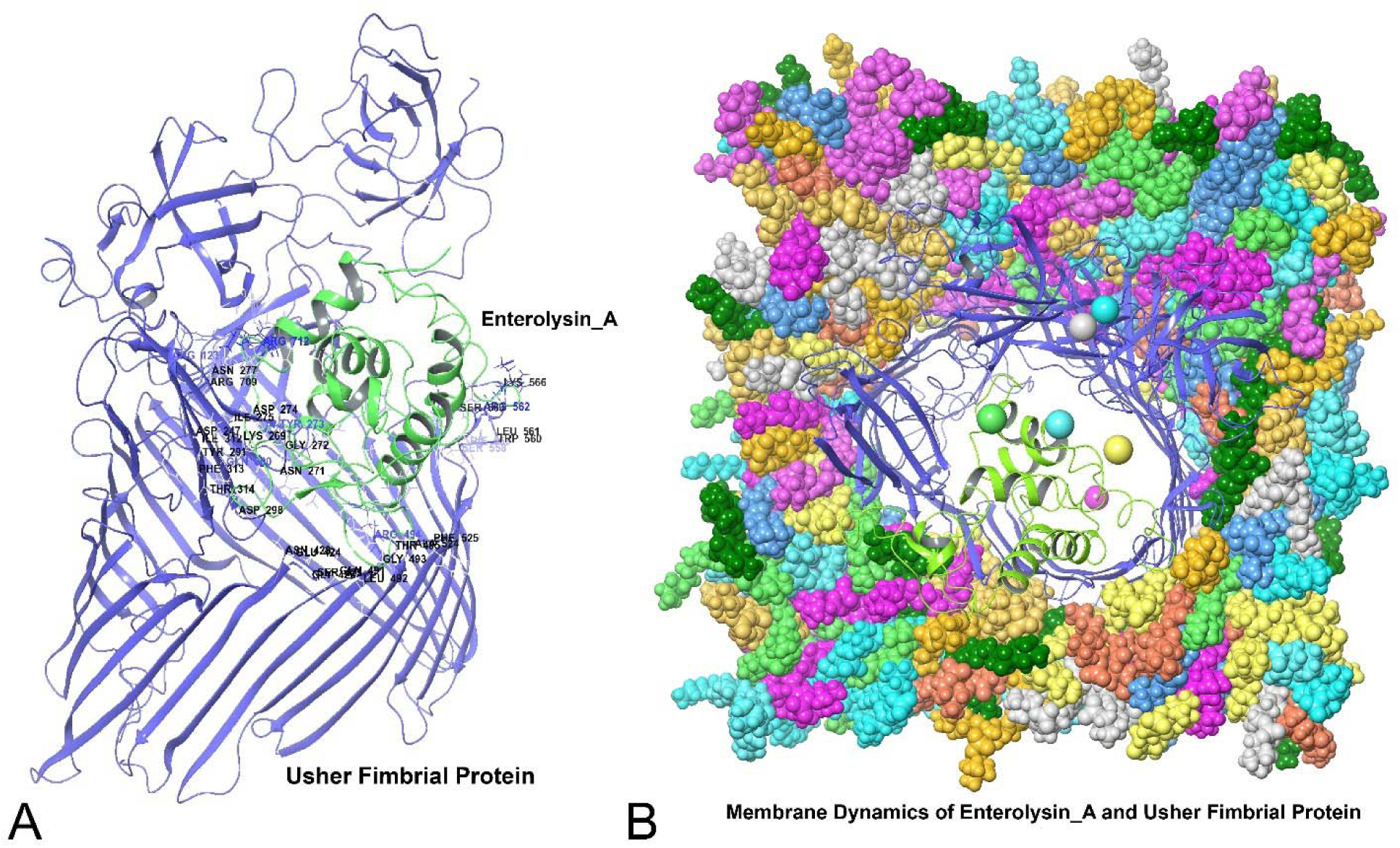
Molecular interactions between bacteriocin compound (e.g., Enterolysin_A) of *E. faecium* MBBL3 and virulence protein (e.g., usher protein) of bovine clinical mastitis pathogens *K. pneumoniae* MBBL2 and *E. coli* MBBL4. (A) Enterolysin_A formed a complex with usher protein in the channel regions of the protein, which indicates potentially impairing its function. (B) Membrane dynamics of Enterolysin_A-usher protein complex, which were found stable throughout the 200 ns dynamics trajectories.

## Discussion

The global rise in multidrug resistance (MDR) bacteria presents a critical clinical health challenge for both human and animal populations, with projections indicating it could result in 10 million human deaths annually by 2050 (Zaghloul and Halfawy 2024). The indiscriminate use of antibiotics in veterinary medicine, particularly for managing mastitis in dairy animals, accelerates the development of antimicrobial resistance in associated bacterial communities, encompassing both pathogenic and commensal species. Mitigating this challenge underscores the urgent need to explore and develop sustainable alternatives to conventional antibiotic therapies. In recent years, probiotics have gained substantial scientific attention as a promising alternative to antibiotics, due to their potential therapeutic applications, antimicrobial properties, and roles in food preservation (Hasnat et al. 2024b; Liu et al. 2023).

In this study, WGS analysis of *E. faecium* MBBL3 revealed a close genetic relationship with other *Enterococcus* strains, along with significant probiotic and antimicrobial properties. The genome of MBBL3 was notably enriched in pathways related to essential carbohydrate metabolism, primary and secondary metabolites, antimicrobial compounds, and a favorable safety profile. These genomic characteristics, combined with its *in-vitro* and *in-silico* efficacy in inhibiting the growth of key mastitis pathogens, position MBBL3 as a safe probiotic and a promising candidate for bioactive natural therapeutics. Importantly, *E. faecium* MBBL3 genome showed a close genetic relationship with the potential probiotic strain *E. lactis* DH9003, isolated from goat milk (Yang et al. 2023). Furthermore, OrthoANI, pangenome, and orthogroup analyses revealed that the *E. faecium* MBBL3 strain shares significant genetic similarity with other *Enterococcus* probiotic strains namely *E. faecium* HY07 (Duan et al. 2019), *E. faecium* BIOPOP- 3WT and BIOPOP-3ALE (Min et al. 2020), and *E. lactis* DH9003 (Yang et al. 2023). Despite originating from different sources *viz*. *E. faecium* MBBL3 from milk and *E. faecium* HY07 from Chinese sausages, the two genomes shared extensive conserved regions, with some genes exhibiting inversions or positional rearrangements within their respective chromosomes. These findings underscore the conserved nature of open reading frames in the reference genome while suggesting that observed variations may be attributable to differences in genome assembly quality. Biochemical profiling of MBBL3 demonstrated its strong fermentation ability with a variety of sugars, alongside significant enzymatic activities. Notably, MBBL3 efficiently fermented essential carbohydrates such as maltose, mannose, sucrose, trehalose, N- acetylglucosamine, and arginine (**Table 1**). Carbohydrate utilization is a fundamental trait of probiotics, vital for their growth, metabolic functions, colonization of the gastrointestinal tract, and overall efficacy. The MBBL3 genome contained 76 CAZymes distributed across five CAZy families, with some exhibiting antimicrobial properties. The predicted CAZymes can contribute to antimicrobial defense by targeting bacterial and fungal cell walls (Onyango et al. 2021; Tathode et al. 2024). For instance, GH73 hydrolyzes peptidoglycan and causes bacterial lysis (Rahman et al. 2024b), while GH18 and CBM50 degrade chitin, and exhibit antifungal activity (Costa et al. 2020; Zhou et al. 2024). CE4 hydrolyzes ester bonds, and increase bacterial and fungal cell walls susceptibility to enzymatic degradation (Ameri et al. 2022). AA10 enzyme, also known as lytic polysaccharide monooxygenases, degrade polysaccharides like chitin and cellulose, enhancing the breakdown of bacterial and fungal cell walls and contributing to antimicrobial and antifungal activity (Yao et al. 2023). AA10 families are also known for their ability to disrupt biofilm matrix polysaccharides, potentially facilitating antibacterial action (Yao et al. 2023). In this study, the comprehensive KEGG pathway analysis of MBBL3 provided valuable insights into its metabolic capabilities, highlighting its efficient energy production in both aerobic and anaerobic conditions. The genome also encodes essential amino acid metabolism pathways and bile salt hydrolase activity, supporting its probiotic potential and survival in the gastrointestinal tract. Furthermore, stress-resistance genes for acid, heat, and cold tolerance enhance its adaptability (Rahman et al. 2024b). These genomic features, along with surface adhesion factors, position MBBL3 as a promising probiotic candidate, demonstrating its metabolic versatility and robustness for potential therapeutic applications in human health.

One of the key findings of this study is the prediction of both primary and secondary metabolites in the MBBL3 genome, which underscores its capacity for producing bioactive compounds with potential antimicrobial properties and probiotic benefits. These include the PFOR II pathway, gallic acid metabolism, and the conversion of arginine to hydrogen carbonate. These metabolic capabilities contribute to several health benefits, including antioxidant, antimicrobial, and anti-inflammatory activities (Rahman et al. 2024b). The genome contained secondary metabolite regions responsible for the biosynthesis of 2,4-DAPG, aborycin, enterocin NKR-5-3B, and sodorifen, highlighting its potential to produce diverse bioactive compounds with antimicrobial and signaling properties. In previous studies, 2,4-DAPG exhibited broad-spectrum antimicrobial activity against human pathogens such as *S. aureus* and *E. coli*, disrupting bacterial cell membranes and inhibiting biofilm metabolic activity, thereby reducing resistance to antimicrobial treatments (Julian et al. 2020; Zhong et al. 2023). Aborycin demonstrated antimicrobial efficacy against *S. aureus*, *E. faecalis*, and *Enterococcus gallinarum*, and was also identified as an anti-HIV metabolite (Li et al. 2023; Shao et al. 2019). Enterocin NKR-5-3B exhibited potent antimicrobial activity against *S. aureus*, *B. cereus*, *E. coli*, *Listeria monocytogenes*, and *Salmonella enterica*, and remained stable across various pH conditions (Kohei et al. 2015; Wang et al. 2023). Sodorifen was recognized for its antifungal properties and may have contributed to ecological bacterial interactions, further enhancing its antimicrobial activity (Bustamante et al. 2022). Additionally, the genome contains BBGCs, including those for sactipeptides, Enterolysin_A and UviB, suggesting MBBL3 can produce antimicrobial peptides. The ABC transporter proteins predicted in the sactipeptides are expected to serve a dual function, primarily involved in multidrug resistance and the import of ATP-binding proteins (Zaghloul and Halfawy 2024). Enterolysin A disrupts the cell walls of Gram-positive bacteria, leading to cell lysis, and has already shown promise as an antimicrobial agent, particularly for combating MDR bacterial infections (Almeida-Santos et al. 2021). On the other hand, UviB, a novel bacteriocin, plays a crucial role in bacteriophage-host interactions by regulating bacterial cell wall degradation (Hong et al. 2022). We identified only two ARGs, corroborating the AST findings, and one VFG (*acm*) for cellular adhesion. The MBBL3 genome harbors three CRISPR regions, with two Cas clusters identified as MGEs. The absence of ARGs and VFGs within these MGEs suggests a favorable safety profile for therapeutic applications. Furthermore, absence of hemolytic activity makes MBBL3 a safe and promising candidate for therapeutic development and biotechnological applications.

A notable observation from this research was that *E. faecium* MBBL3 effectively inhibited the growth of *K. pneumoniae* (*Kp* MBBL2) and *E. coli* (*Ec* MBBL4), two major mastitis pathogens, in *in-vitro* assays. Additionally, this study elucidated a potential molecular mechanism underlying this inhibition in *in-silico*. Our analysis showed that Enterolysin A strongly interacts with the outer membrane usher protein, a key component in pili assembly and transport, processes vital for bacterial adhesion, biofilm formation, and colonization, essential factors in the virulence of Gram-negative bacteria such as *K. pneumoniae* and *E. coli* (Gato et al. 2020; Olson et al. 2024). The binding of Enterolysin A to the usher protein reveals a potential mechanism through which MBBL3 exerts its inhibitory effects, likely disrupting adhesion, biofilm formation, and colonization, thereby inhibiting the survival and proliferation of *K. pneumoniae* and *E. coli*. This robust interaction is likely the key driver behind MBBL3’s antimicrobial activity, as demonstrated in our study.

In summary, this study offers a comprehensive bioinformatics analysis of *E. faecium* MBBL3, comparing its genome with other *Enterococcus* strains and evaluating its probiotic potential and antimicrobial efficacy against mastitis-causing pathogens. Non-hemolytic in nature, extensive ranges of carbohydrates fermentation capability, along with possesses genes linked to adaptation and survivability to various environment including gut of the host. Additionally, it’s CAZymes, primary and secondary metabolites, bacteriocin gene clusters -related to antimicrobial highlighting its potential antimicrobial capabilities. *In-vitro* tests and *In-silico* genomic analysis with *Enterolysin_A* of *E. faecium* MBBL3 further supported its strong inhibitory effects against mastitis pathogens *Kp* MBBL2 and *Ec* MBBL4. These findings establish *E. faecium* MBBL3 as a promising candidate for biotherapeutics targeting mastitis, with additional potential for applications in biopreservation and bioremediation. The genomic insights provide a foundation for exploring its probiotic and therapeutic roles in animal models and interactions with gut microbiota, supporting the development of probiotic-based strategies to manage mastitis and improve dairy herd health and productivity.

## Authors’ contributions

N.S., M.M.R. and M.N.H. conceived and designed the study. N.S., M.M.R. and S.H. curated, analyzed and visualized data, and wrote original draft. A.N.M.A.R., A.K.T., M.R.K., Z.C.D., T.I. and M.N.H. critically reviewed and edited the manuscript. All authors have read and approved the final manuscript.

## Data Availability

The whole genome sequence of *Enterococcus faecium* MBBL3 has been archived in the NCBI GenBank and the NCBI Sequence Read Archive (SRA) under BioProject accession PRJNA1065156 and SRA accession SRR27558663. The version reported in this study is identified as version JAZIFO000000000.1.

## Declarations Ethics approval

This study received ethical approval from the Animal Research Ethics Committee (AREC) of Bangabandhu Sheikh Mujibur Rahman Agricultural University, Bangladesh, under reference number FVMAS/AREC/2024/6755, granted on January 20, 2024. Furthermore, this article does not contain any studies with human participants or animals performed by any of the authors.

