## Supplementary material for "*Enterococcus faecium* MBBL3 Exhibits Promising Probiotic Potential and Antimicrobial Efficacy Against Bovine Mastitis-Associated *Escherichia coli and Klebsiella pneumoniae*": Table S1-S8

**Supplementary Tables**

**Table S1.** Accession ID, isolation source, and origin of 14 E. faecium and one E. lactis strains.

| **Sl No.** | **Accession ID** | **Strain** | **Isolation source** | **Country** |
| --- | --- | --- | --- | --- |
| 1 | GCA_036620595.1 | *E. faecium* MBBL3 | Milk | Bangladesh |
| 2 | CP072707.1 | *E. faecium* VB3378 | Blood | India |
| 3 | JRGX00000000.1 | *E. faecium* L-3 | Probiotic preparation | Russia |
| 4 | CP079880.1 | *E. lactis* CX 2-6_2 | Soil | China |
| 5 | CP043484.1 | *E. faecium* DMEA02 | Meju, fermented soybean | South Korea |
| 6 | CP129887.1 | *E. lactis* SU-B46 | Milk | Argentina |
| 7 | CP040878.1 | *E. faecium* HB-1 | Feces | South Korea |
| 8 | CP035136.1 | *E. faecium* SRCM103341 | Food | South Korea |
| 9 | CP033376.1 | *E. faecium* Gr17 | Fermented whole fish product suan yu | China |
| 10 | CP059793.1 | *E. faecium* BP3378 | Blood | India |
| 11 | CP032308.1 | *E. faecium* HY07 | Sausages | China |
| 12 | CP050650.1 | *E. faecium* BIOPOP-3WT | Fermented dairy products | South Korea |
| 13 | CP050648.1 | *E. faecium* BIOPOP-3ALE | Fermented dairy products | South Korea |
| 14 | CP084248.1 | *E. lactis* DH9003 | Goat milk | China |
| 15 | CP045602.1 | *E. faecium* TK-P5D | Probiotic products | China |

**Table S2.** Essential and virulence proteins of *K. pneumoniae* MBBL2 and *E. coli* MBBL4

| Sl. No. | *K. pneumoniae* MBBL2 | *E. coli* MMBL4 |
| --- | --- | --- |
| 1 | MFH5886825.1 nitrate reductase subunit alpha [Klebsiella pneumoniae] | MFH5812639.1 fimbrial biogenesis usher protein [Escherichia coli] |
| 2 | MFH5887007.1 multidrug efflux RND transporter permease subunit KexD | MFH5812680.1 tyrosine-protein kinase [Escherichia coli] |
| 3 | MFH5887164.1 nitrate reductase subunit alpha [Klebsiella pneumoniae] | MFH5812793.1 HilA family transcriptional regulator YgeH [Escherichia |
| 4 | MFH5887197.1 VasL domain-containing protein [Klebsiella pneumoniae] | MFH5813680.1 NADP-dependent phosphogluconate dehydrogenase |
| 5 | MFH5887201.1 type VI secretion system baseplate subunit TssF | MFH5813718.1 efflux RND transporter permease AcrB [Escherichia coli] |
| 6 | MFH5887218.1 DUF2345 domain-containing protein [Klebsiella | MFH5813815.1 carbamoyl-phosphate synthase large subunit [Escherichia |
| 7 | MFH5887219.1 OmpA family protein [Klebsiella pneumoniae] | MFH5813888.1 type II secretion system protein GspE [Escherichia |
| 8 | MFH5887221.1 type VI secretion system baseplate subunit TssK | MFH5814193.1 two-component system sensor histidine kinase PhoQ |
| 9 | MFH5887253.1 TonB-dependent siderophore receptor [Klebsiella | MFH5814239.1 flagellar hook-associated protein FlgK [Escherichia |
| 10 | MFH5887396.1 ATP-dependent chaperone ClpB [Klebsiella pneumoniae] | MFH5814315.1 magnesium-translocating P-type ATPase [Escherichia |
| 11 | MFH5887486.1 multidrug efflux RND transporter permease subunit OqxB | MFH5814409.1 glucose-6-phosphate isomerase [Escherichia coli] |
| 12 | MFH5887787.1 type 3 fimbria usher protein MrkC [Klebsiella | MFH5814424.1 SopA family protein [Escherichia coli] |
| 13 | MFH5887800.1 fimbrial biogenesis usher protein [Klebsiella | MFH5814450.1 T3SS effector pentapeptide repeat protein EspX5 |
| 14 | MFH5887804.1 EAL domain-containing protein [Klebsiella pneumoniae] | MFH5814901.1 (2,3-dihydroxybenzoyl)adenylate synthase EntE |
| 15 | MFH5888036.1 type VI secretion system baseplate subunit TssF | MFH5814989.1 phosphoethanolamine transferase CptA [Escherichia coli] |
| 16 | MFH5888037.1 type VI secretion system protein TssA [Klebsiella | MFH5814909.1 enterobactin non-ribosomal peptide synthetase EntF |
| 17 | MFH5888038.1 ImcF-related family protein [Klebsiella pneumoniae] | MFH5814998.1 flagellar protein export ATPase FliI [Escherichia coli] |
| 18 | MFH5888041.1 T6SS phospholipase effector Tle1-like catalytic | MFH5814912.1 siderophore enterobactin receptor FepA [Escherichia |
| 19 | MFH5888046.1 type VI secretion system Vgr family protein [Klebsiella | MFH5815533.1 ATP-dependent chaperone ClpB [Escherichia coli] |
| 20 | MFH5888047.1 type VI secretion system ATPase TssH [Klebsiella | MFH5815682.1 flagellar biosynthesis protein FlhA [Escherichia coli] |
| 21 | MFH5888049.1 OmpA family protein [Klebsiella pneumoniae] | MFH5814995.1 flagellar basal-body MS-ring/collar protein FliF |
| 22 | MFH5888051.1 type VI secretion system baseplate subunit TssK | MFH5815691.1 chemotaxis protein CheA [Escherichia coli] |
| 23 | MFH5888052.1 type VI secretion system contractile sheath large | MFH5815472.1 multidrug efflux RND transporter permease subunit AcrF |
| 24 | MFH5888139.1 TonB-dependent siderophore receptor [Klebsiella | MFH5816393.1 allantoinase AllB [Escherichia coli] |
| 25 | MFH5888920.1 (2,3-dihydroxybenzoyl)adenylate synthase EntE | MFH5816474.1 nitrate reductase subunit alpha [Escherichia coli] |
| 26 | MFH5889323.1 CS1-pili formation C-terminal domain-containing protein | MFH5815689.1 methyl-accepting chemotaxis protein II [Escherichia |
| 27 | MFH5889324.1 fimbrial adhesin EcpD [Klebsiella pneumoniae] | MFH5816581.1 nitrate reductase Z subunit alpha [Escherichia coli] |
| 28 | MFH5889477.1 multidrug efflux RND transporter permease subunit AcrB | MFH5815723.1 flagellar filament capping protein FliD [Escherichia |
| 29 | MFH5889620.1 magnesium-translocating P-type ATPase [Klebsiella | MFH5816649.1 rhs element protein RhsD, partial [Escherichia coli] |
| 30 | MFH5890488.1 capsule assembly Wzi family protein [Klebsiella | MFH5816652.1 rhs element protein RhsB, partial [Escherichia coli] |
| 31 | MFH5890790.1 glucose-6-phosphate isomerase [Klebsiella pneumoniae] | MFH5816030.1 beta strand repeat-containing protein, partial [Escherichia coli] |
| 32 | MFH5890826.1 TonB-dependent siderophore receptor [Klebsiella | MFH5816484.1 multidrug efflux pump RND permease MdtF [Escherichia coli] |
| 33 | MFH5891186.1 outer membrane usher protein [Klebsiella pneumoniae] |  |
| 34 | MFH5891259.1 efflux RND transporter permease subunit [Klebsiella] |  |

**Table S3.** Complete KEGG metabolic pathway modules in the *E. faecium* MBBL3 genome.

| Pathway modules | Category | Complete pathway modules | KEGG Module ID |
| --- | --- | --- | --- |
| Carbohydrate metabolism | Glycolysis / Gluconeogenesis | Glycolysis (Embden-Meyerhof pathway) | [M00001](https://www.kegg.jp/kegg-bin/show_module?17320786674190328/M00001.args.multi) |
|  |  | Glycolysis, core module involving three-carbon compounds | [M00002](https://www.kegg.jp/kegg-bin/show_module?17320786674190328/M00002.args.multi) |
|  |  | Gluconeogenesis | [M00003](https://www.kegg.jp/kegg-bin/show_module?17320786674190328/M00003.args.multi) |
|  |  | Pyruvate oxidation | [M00307](https://www.kegg.jp/kegg-bin/show_module?17320786674190328/M00307.args.multi) |
|  | Pentose phosphate pathway | Pentose phosphate pathway, oxidative phase | [M00006](https://www.kegg.jp/kegg-bin/show_module?17320786674190328/M00006.args.multi) |
|  |  | PRPP biosynthesis | [M00005](https://www.kegg.jp/kegg-bin/show_module?17320786674190328/M00005.args.multi) |
| Other carbohydrate metabolism | Galactose metabolism | Galactose degradation, Leloir pathway | [M00632](https://www.kegg.jp/kegg-bin/show_module?17320786674190328/M00632.args.multi) |
|  | Ascorbate and aldarate metabolism | Ascorbate degradation | [M00550](https://www.kegg.jp/kegg-bin/show_module?17320786674190328/M00550.args.multi) |
|  | Amino sugar and nucleotide sugar metabolism | Nucleotide sugar biosynthesis | [M00549](https://www.kegg.jp/kegg-bin/show_module?17320786674190328/M00549.args.multi) |
|  |  | Nucleotide sugar biosynthesis | [M00554](https://www.kegg.jp/kegg-bin/show_module?17320786674190328/M00554.args.multi) |
|  |  | UDP-N-acetyl-D-glucosamine biosynthesis | [M00909](https://www.kegg.jp/kegg-bin/show_module?17320786674190328/M00909.args.multi) |
| Energy metabolism | Carbon fixation | Phosphate acetyltransferase-acetate kinase pathway | [M00579](https://www.kegg.jp/kegg-bin/show_module?17320786674190328/M00579.args.multi) |
|  | ATP synthesis | F-type ATPase, prokaryotes and chloroplasts | [M00157](https://www.kegg.jp/kegg-bin/show_module?17320786674190328/M00157.args.multi) |
|  |  | V/A-type ATPase, prokaryotes | [M00159](https://www.kegg.jp/kegg-bin/show_module?17320786674190328/M00159.args.multi) |
| Lipid metabolism | Fatty acid metabolism | Fatty acid biosynthesis, initiation | [M00082](https://www.kegg.jp/kegg-bin/show_module?17320786674190328/M00082.args.multi) |
|  |  | Fatty acid biosynthesis, elongation | [M00083](https://www.kegg.jp/kegg-bin/show_module?17320786674190328/M00083.args.multi) |
| Nucleotide metabolism | Purine metabolism | De novo purine biosynthesis | [M00048](https://www.kegg.jp/kegg-bin/show_module?17320786674190328/M00048.args.multi) |
|  |  | Adenine ribonucleotide biosynthesis | [M00049](https://www.kegg.jp/kegg-bin/show_module?17320786674190328/M00049.args.multi) |
|  |  | Guanine ribonucleotide biosynthesis | [M00050](https://www.kegg.jp/kegg-bin/show_module?17320786674190328/M00050.args.multi) |
|  |  | Deoxyribonucleotide biosynthesis | [M00053](https://www.kegg.jp/kegg-bin/show_module?17320786674190328/M00053.args.multi) |
|  | Pyrimidine metabolism | Pyrimidine ribonucleotide biosynthesis | [M00052](https://www.kegg.jp/kegg-bin/show_module?17320786674190328/M00052.args.multi) |
|  |  | Pyrimidine deoxyribonucleotide biosynthesis | [M00938](https://www.kegg.jp/kegg-bin/show_module?17320786674190328/M00938.args.multi) |
| Amino acid metabolism | Cysteine and methionine metabolism | Cysteine biosynthesis | [M00021](https://www.kegg.jp/kegg-bin/show_module?17320786674190328/M00021.args.multi) |
|  | Arginine and proline metabolism | Proline biosynthesis | [M00015](https://www.kegg.jp/kegg-bin/show_module?17320786674190328/M00015.args.multi) |
| Metabolism of cofactors and vitamins | Cofactor and vitamin metabolism | Coenzyme A biosynthesis | [M00120](https://www.kegg.jp/kegg-bin/show_module?17320786674190328/M00120.args.multi) |
|  |  | C1-unit interconversion | [M00140](https://www.kegg.jp/kegg-bin/show_module?17320786674190328/M00140.args.multi) |
| Biosynthesis of terpenoids and polyketides | Terpenoid backbone biosynthesis | C10-C20 isoprenoid biosynthesis, bacteria | [M00364](https://www.kegg.jp/kegg-bin/show_module?17320786674190328/M00364.args.multi) |
|  | Polyketide sugar unit biosynthesis | dTDP-L-rhamnose biosynthesis | [M00793](https://www.kegg.jp/kegg-bin/show_module?17320786674190328/M00793.args.multi) |

**Table S4.** Prediction of primary and secondary metabolite biosynthetic gene clusters in E. faecium MBBL3.

| Region | Type | From | To | Most similar known clusters |
| --- | --- | --- | --- | --- |
| Primary metabolite region 1 | Pyruvate2acetate-formate | 6,324 | 29,560 | Pyruvate to acetate and formate *C. acetobutylicum*, PFL_acetate  (100% similarity) |
| Primary metabolite region 2 | PFOR_II_pathway | 1 | 15,753 | PFOR II pathway *B. thetaiotaomicroni*, PFORII  (100% similarity) |
| Primary metabolite region 3 | gallic_acid_met | 29,167 | 51,273 | Gallic acid degradation *B. sp.* KLE, GALL  (100% similarity) |
| Primary metabolite region 4 | Arginine2_Hcarbonate | 1 | 21,707 | Arginine to hydrogen carbonate *P. aeruginosa*, ARG (100% similarity) |
| Secondary metabolite region 1 | T3PKS | 77,070 | 118,224 | - |
| Secondary metabolite region 2 | cyclic-lactone-autoinducer | 1 | 15,509 | - |
| Secondary metabolite region 3 | cyclic-lactone-autoinducer | 17,186 | 31,091 | - |
| Secondary metabolite region 4 | cyclic-lactone-autoinducer | 1 | 13,147 | - |

**Table S5.** Minimum Information about a Biosynthetic Gene cluster (MIBiG) comparison similarity score (region to region analysis) of secondary metabolite biosynthesis gene clusters of *E. faecium* MBBL3.

| **Secondary metabolites** | **Reference** | **Similarity score** | **Type** | **Compound(s)** | **Organism** |
| --- | --- | --- | --- | --- | --- |
| T3PKS | [BGC0000867](https://mibig.secondarymetabolites.org/repository/BGC0000867/index.html#r1c1) | 0.28 | Other | polyhydroxyalkanoic acids | *Ectothiorhodospira shaposhnikovii* |
|  | [BGC0000866](https://mibig.secondarymetabolites.org/repository/BGC0000866/index.html#r1c1) | 0.27 | Other | polyhydroxyalkanoate | *Burkholderia* sp. DSM 9242 |
|  | [BGC0000280](https://mibig.secondarymetabolites.org/repository/BGC0000280/index.html#r1c1) | 0.27 | Polyketide | 2,4-diacetylphloroglucinol | *Pseudomonas fluorescens* |
|  | [BGC0000281](https://mibig.secondarymetabolites.org/repository/BGC0000281/index.html#r1c1) | 0.25 | Polyketide | 2,4-diacetylphloroglucinol | *Pseudomonas fluorescens* |
|  | [BGC0000286](https://mibig.secondarymetabolites.org/repository/BGC0000286/index.html#r1c1) | 0.23 | Polyketide | viguiepinol | *Streptomyces* sp*.* KO-3988 |
|  | [BGC0002561](https://mibig.secondarymetabolites.org/repository/BGC0002561/index.html#r1c1) | 0.20 | Alkaloid | phenazine SA, phenazine SB, phenazine SC | *Streptomyces* sp. |
|  | [BGC0000255](https://mibig.secondarymetabolites.org/repository/BGC0000255/index.html#r1c1) | 0.20 | Polyketide | pederin | Uncultured bacterium |
|  | [BGC0000205](https://mibig.secondarymetabolites.org/repository/BGC0000205/index.html#r1c1) | 0.16 | Polyketide | bryostatin | *Candidatus Endobugula sertula* |
|  | [BGC0001080](https://mibig.secondarymetabolites.org/repository/BGC0001080/index.html#r1c1) | 0.14 | Other (Phenazine) | endophenazine A, endophenazine B | *Streptomyces anulatus* |
|  | [BGC0002584](https://mibig.secondarymetabolites.org/repository/BGC0002584/index.html#r1c1) | 0.13 | RiPP | azolemycin B, azolemycin A, azolemycin D, azolemycin C | *Streptomyces* sp. FXJ1.264 |
| Cyclic-lactone-autoinducer | [BGC0002285](https://mibig.secondarymetabolites.org/repository/BGC0002285/index.html#r1c1) | 0.20 | RiPP | aborycin | *Streptomyces* sp. ZS0098 |
|  | [BGC0001291](https://mibig.secondarymetabolites.org/repository/BGC0001291/index.html#r1c1) | 0.19 | RiPP | enterocin NKR-5-3B | *Enterococcus faecium* |
|  | [BGC0000540](https://mibig.secondarymetabolites.org/repository/BGC0000540/index.html#r1c1) | 0.12 | RiPP | paenibacillin | *Paenibacillus polymyxa* OSY-DF |
|  | [BGC0002585](https://mibig.secondarymetabolites.org/repository/BGC0002585/index.html#r1c1) | 0.09 | Other | ubericin K | *Streptococcus uberis* |
|  | [BGC0002579](https://mibig.secondarymetabolites.org/repository/BGC0002579/index.html#r1c1) | 0.09 | RiPP | carnobacteriocin XY | *Carnobacterium maltaromaticum* |
|  | [BGC0000624](https://mibig.secondarymetabolites.org/repository/BGC0000624/index.html#r1c1) | 0.09 | RiPP | salivaricin CRL1328 α peptide, salivaricin CRL1328 β peptide | *Lactobacillus salivarius* |
|  | [BGC0002667](https://mibig.secondarymetabolites.org/repository/BGC0002667/index.html#r1c1) | 0.08 | RiPP | estericin A | *Clostridium estertheticum* |
|  | [BGC0001510](https://mibig.secondarymetabolites.org/repository/BGC0001510/index.html#r1c1) | 0.07 | Other | anisomycin | *Streptomyces hygrospinosus* |
|  | [BGC0000804](https://mibig.secondarymetabolites.org/repository/BGC0000804/index.html#r1c1) | 0.05 | Saccharide | acarviostatin I03, acarviostatin II03, acarviostatin III03, acarviostatin IV03 | *Streptomyces coelicoflavus* ZG0656 |
|  | [BGC0001728](https://mibig.secondarymetabolites.org/repository/BGC0001728/index.html#r1c1) | 0.03 | NRP, Polyketide | paenilipoheptin | *Paenibacillus polymyxa* E681 |
| Cyclic-lactone-autoinducer | [BGC0001291](https://mibig.secondarymetabolites.org/repository/BGC0001291/index.html#r1c1) | 0.18 | RiPP | enterocin NKR-5-3B | *Enterococcus faecium* |
|  | [BGC0002579](https://mibig.secondarymetabolites.org/repository/BGC0002579/index.html#r1c1) | 0.15 | RiPP | carnobacteriocin XY | *Carnobacterium maltaromaticum* |
|  | [BGC0001388](https://mibig.secondarymetabolites.org/repository/BGC0001388/index.html#r1c1) | 0.11 | RiPP | gassericin E | *Lactobacillus gasseri* |
|  | [BGC0000619](https://mibig.secondarymetabolites.org/repository/BGC0000619/index.html#r1c1) | 0.11 | RiPP | gassericin T | *Lactobacillus gasseri* |
|  | [BGC0001602](https://mibig.secondarymetabolites.org/repository/BGC0001602/index.html#r1c1) | 0.11 | RiPP | gassericin T | *Lactobacillus gasseri* |
|  | [BGC0000540](https://mibig.secondarymetabolites.org/repository/BGC0000540/index.html#r1c1) | 0.10 | RiPP | paenibacillin | *Paenibacillus polymyxa* OSY-DF |
|  | [BGC0002667](https://mibig.secondarymetabolites.org/repository/BGC0002667/index.html#r1c1) | 0.09 | RiPP | estericin A | *Clostridium estertheticum* |
|  | [BGC0000607](https://mibig.secondarymetabolites.org/repository/BGC0000607/index.html#r1c1) | 0.09 | RiPP | micrococcin P1 | *Macrococcus caseolyticus* |
|  | [BGC0000522](https://mibig.secondarymetabolites.org/repository/BGC0000522/index.html#r1c1) | 0.09 | RiPP | lacticin Z | *Lactococcus lactis* |
| Cyclic-lactone-autoinducer | [BGC0001361](https://mibig.secondarymetabolites.org/repository/BGC0001361/index.html#r1c1) | 0.26 | Terpene | sodorifen | *Serratia plymuthica* 4Rx13 |
|  | [BGC0002283](https://mibig.secondarymetabolites.org/repository/BGC0002283/index.html#r1c1) | 0.26 | Terpene | sodorifen | *Serratia plymuthica* |
|  | [BGC0001291](https://mibig.secondarymetabolites.org/repository/BGC0001291/index.html#r1c1) | 0.18 | RiPP | enterocin NKR-5-3B | *Enterococcus faecium* |
|  | [BGC0002579](https://mibig.secondarymetabolites.org/repository/BGC0002579/index.html#r1c1) | 0.15 | RiPP | carnobacteriocin XY | *Carnobacterium maltaromaticum* |
|  | [BGC0001594](https://mibig.secondarymetabolites.org/repository/BGC0001594/index.html#r1c1) | 0.15 | Alkaloid | fischerindole L | *Fischerella muscicola* UTEX 1829 |
|  | [BGC0002585](https://mibig.secondarymetabolites.org/repository/BGC0002585/index.html#r1c1) | 0.14 | Other | ubericin K | *Streptococcus uberis* |
|  | [BGC0000668](https://mibig.secondarymetabolites.org/repository/BGC0000668/index.html#r1c1) | 0.14 | Terpene, Alkaloid | 12-epi-hapalindole C isonitrile, 12-epi-hapalindole E, 12-epi-fischerindole U isonitrile, fischerindole L, 12-epi-fischerindole I isonitrile, welwitindolinone A isonitrile, welwitindolinone B isothiocyanate, welwitindolinone C isothiocyanate, N-methylwelwitindolinone C isothiocyanate, N-methylwelwitinsolinone C isonitrile, 3-epi-welwitindolinone B isothiocyanate, 3-(Z-2'-isocyanoethenyl)-indole | *Hapalosiphon welwitschii* UTEX B 1830 |
|  | [BGC0001501](https://mibig.secondarymetabolites.org/repository/BGC0001501/index.html#r1c1) | 0.13 | Alkaloid | ambiguine P | *Fischerella* sp. TAU |
|  | [BGC0001126](https://mibig.secondarymetabolites.org/repository/BGC0001126/index.html#r1c1) | 0.12 | Terpene, Alkaloid | 12-epi-hapalindole J isonitrile, ambiguine A isonitrile, ambiguine B isonitrile, ambiguine C isonitrile, ambiguine D isonitrile, ambiguine E isonitrile, ambiguine K isonitrile, ambiguine L isonitrile, ambiguine I isonitrile, ambiguine J isonitrile | *Fischerella ambigua* UTEX 1903 |
|  | [BGC0001612](https://mibig.secondarymetabolites.org/repository/BGC0001612/index.html#r1c1) | 0.12 | Alkaloid | ambiguine H isonitrile | *Fischerella ambigua* UTEX 1903 |

**Table S6.** Antibiotic susceptibility results from culture plate: R = Resistance, S = Susceptible, SDD = Susceptible dose-dependent and I = intermediate AST results. The third column in the right column represents zone of inhibition (ZI).

| Antibiotic | AST result | ZI (mm) |
| --- | --- | --- |
| Azithromycin (AZM) | R | 0 |
| Amikacin (AK) | R | 5 |
| Cefepime (FEP) | SDD | 20 |
| Ceftriaxone (CRO) | S | 25 |
| Cefuroxime sodium (CXM) | S | 20 |
| Ciprofloxacin (CIP) | S | 21 |
| Gentamicin (CN) | R | 5 |
| Nalidixic acid (NA) | I | 14 |
| Imipenem (IPM) | S | 22 |
| Meropenem (MEM) | S | 27 |

**Table S7.** Prediction of antibiotic resistance genes in E. faecium MBBL3.

| AMR gene | Start | End | Strand | Drug resistance | Database |
| --- | --- | --- | --- | --- | --- |
| *msr(C)* | 80014 | 81492 | - | Macrolide | CARD |
| *aac(6')-Ii* | 69972 | 70520 | - | Aminoglycoside | CARD |

**Table S8.** Prediction of CRISPER/Cas in E. faecium MBBL3.

| Element | CRISPR Id /  Cas Type | Start | End | Strand |
| --- | --- | --- | --- | --- |
| CRISPER 1 | NZ_JAZIFO010000001.1_1 | 490,921 bp | 491,014 bp | + |
| CRISPER 2 | NZ_JAZIFO010000001.1_2 | 732,796 bp | 733,227 bp | + |
| CRISPER 3 | NZ_JAZIFO010000001.1_3 | 2101,223 bp | 2101,421 | + |
| Cas cluster | General-Class1 | 613,690 | 153,5691 | + |
| Cas cluster | General-Class2 | 726,823 | 732,705 | + |
